# Metabolomics and Microbiomics Insights into the Differential Surface Fouling of Brown Algae

**DOI:** 10.1101/2023.07.14.548367

**Authors:** Ernest Oppong-Danquah, Martina Blümel, Deniz Tasdemir

## Abstract

Marine macroalgae (seaweeds) are key components of marine ecosystems with vital roles in costal habitats. As they release dissolved organic matter and other molecules, seaweeds are under strong settlement pressure by micro- and macro-epibionts. Uncontrolled epibiosis causes surface fouling with detrimental effects on the health and well-being of the organism. Seaweeds control surface epibionts directly by releasing antifouling and antimicrobial metabolites onto their surfaces, and indirectly by recruiting beneficial microorganisms that produce antimicrobial/antifouling metabolites. Three species of the brown algal genus *Fucus, F. vesiculosus* (FV)*, F. serratus* (FS) and *F. distichus* subsp. *evanescens* (FE) form the *Fucus* belt habitat in the Kiel Fjord, Germany. They often co-occur in the same spot but their blades are fouled differently; we observed FE to be the least fouled, and FV to be the most fouled species. This study was designed to investigate the potential factors underlying different fouling intensities on the surfaces of the three co-occurring *Fucus* spp. Their surface metabolomes were analysed by comparative untargeted UPLC-MS/MS based metabolomics to identify marker metabolites influencing the surface fouling. The epiphytic microbial communities of the *Fucus* spp. were also comparatively characterized by high-throughput amplicon sequencing to identify the differences in the surface microbiome of the algae. By employing these omics methods, integrated with multivariate analyses, we identified discriminant metabolites and microbial taxa associated with FE surface, including antimicrobial polar lipids, the fungal genera *Mucor*, *Alternaria*, and bacterial genus *Yoonia-Loktanella*. These taxa have been previously reported to produce antimicrobial and antifouling compounds, suggesting their potential involvement in the fouling resistance (least fouled) observed on the FE surface relative to the co-occurring algae FS and FV. These findings shed light on the surface metabolome and microbiome of *Fucus* spp. and their influence in different fouling intensities and also have implications for the conservation of coastal habitats.

## 1. Introduction

Marine macroalgae (seaweeds) are diverse, multicellular, photosynthetic plant-like organisms, which are ubiquitous in coastal ecosystems. They serve as good model organisms for diverse scientific endeavors, including ecology, natural product discovery and host-microbe interactions in the sea (Creed et al., 1997; Wahl et al., 2012; Buedenbender et al., 2020). Macroalgae are one of the key marine ecosystem engineers, contributing significantly to nitrogen, iodine and carbon biogeochemical cycles (Capistrant-Fossa et al., 2021). They are primary producers and also producers of oxygen, thereby reducing the effect of ocean acidification and deoxygenation (Kolanjinathan et al., 2014; Noisette and Hurd, 2018). Seaweeds provide a direct source of food for various marine animals (El-Manaway and Rashedy, 2022). There is also growing interest in seaweeds as blue carbon ecosystems due to their ability as carbon dioxide sink (Krause-Jensen and Duarte, 2016; Ould and Caldwell, 2022).

Seaweeds typically live in the littoral, euphotic zones of the sea. Three *Fucus* spp. (wracks) namely *F. vesiculosus* (bladder wrack; FV), *F. serratus* (toothed or serrated wrack; FS) and *F. distichus* subsp. *evanescens* (two-headed wrack; FE) occur in parallel on hard substrates in Kiel Fjord (German Baltic Sea). These three-foundation species form the *Fucus* belt habitat and provide nursery ground for numerous species, hence contribute significantly to the ecosystem structure and health. While FV and FS are native to Kiel Fjord, FE is an invasive species, which was accidentally introduced from the Arctic region in the late 20^th^ century (Wikström et al., 2002; Coyer et al., 2002). Similar to any other natural or man-made surfaces submerged in seawater, the thalli of algae are prone to fouling, i.e., colonization by other organisms (epibionts). Microorganisms, especially bacteria, are often the primary macroalgal epibionts (Grinberg et al., 2019). The bacteria secrete exopolymers, which (ir)reversibly attach them to the surface (substrata), with subsequent aggregation leading to a thin bacterial biofilm (Monds and O’Toole, 2009). This biofilm forms the basis for further settlement of other microfoulers, such as protozoans, flagellates, fungi, and macrofoulers, e.g., bryozoans or barnacles (Busetti et al., 2017)). Estimated microbial densities in one milliliter of seawater is reported to be 10^3^ fungal cells, 10^6^ bacterial cells and 10^2^ larvae and spores of macrofoulers, depending on season and location (Harder, 2009), increasing the fouling pressure on marine organisms.

Macroalgae have evolved innate physical and chemical defense mechanisms to control epibiosis. Physical defense mechanisms include shedding of surface tissue (epithallus sloughing) as observed in red coralline algae (Littler and Littler, 1999), the brown alga *Ascophyllum nodosum* (Filion-Myklebust, 1981), and the green alga *Ulva intestinalis* (McArthur and Moss, 1977; da Gama et al., 2014). Chemical defense of the macroalgal host involves production and release of oxygen radicals (oxidative bursts) in response to chemical signals from epibionts on the surface (Potin, 2008). Furthermore, macroalgae produce diverse metabolites and release them to their surfaces to actively shape their epibiomes and modulate settlement of colonizers (Givskov et al., 1996; Potin, 2008; Plouguerné et al., 2010). One example for such metabolite is the brominated heptanone, produced and stored in the surface cells of the red alga *Bonnemaisonia hamifera* with antifouling activity (Nylund et al., 2008). The diterpene bromosphaerol and its congeners, isolated from the red alga *Sphaerococcus coronopifolius,* also inhibit the fouling of potential marine bacterial and microalgae epibionts (Quémener et al., 2022).

Macroalgae release dissolved organic matter and chemicals such as sugars and amino acids to attract beneficial microbes onto their surfaces, thereby increasing the selectivity of epibiosis (Harder, 2009; da Gama et al., 2014). In return, some epibionts functionally modulate the performance, elasticity and health of the algal host by supplying essential nutrients and metabolites (Egan et al., 2013). As an example, the bacterium *Mesorhizobium loti* supplies vitamin B12 to the green alga *Lobomonas rostrata* (Croft et al., 2005; Grant et al., 2014), while other epibionts release bioactive compounds, which prevent the settlement of pathogens and other colonizers (Soman et al., 1999; Noorjahan et al., 2022). On the contrary, some epibionts may facilitate further colonization of other prokaryotes and eukaryotes or even increase susceptibility of the host to grazers (Egan et al., 2008; Corrigan et al., 2023). For example, the biofilm-forming *Vibrio anguillarum* enhance the settlement of *Ulva* zoospores, while bryozoan *Membranipora membranacea* on the red seaweed *Cryptonemia seminervis* altered its susceptibility to sea urchin and amphipod grazing (Wheeler et al., 2006; da Gama et al., 2014). These examples illustrate the complex and dynamic nature of interactions between the seaweed host and its epibiome, largely defined on the host surface. Extensive epibiosis may lead to increased weight and surface roughness, consequently increasing the deposition of particulate materials on the algal host surface (Harder, 2009). This further reduces the surface irradiance and oxygen intake leading to poor photosynthesis, stunted growth and eventually promotes algal diseases. Like all macroalgae, *Fucus* spp. also have the capacity to modulate their surface epibionts. The carotenoid fucoxanthin and polyphenol phlorotannins are some examples of known metabolites released onto the surfaces of *Fucus* spp. to control fouling (Honkanen and Jormalainen, 2005; Saha et al., 2011).

In a previous study, we analyzed the surface epibiome and the spatial distribution of the surface metabolome of Kiel Fjord FV by LC-MS/MS and Desorption Electrospray Ionization Imaging Mass Spectrometry (DESI-IMS) (Parrot et al., 2019). Following this initial study, we observed that FV was more intensely fouled compared to the other two co-occurring *Fucus* spp., FS (less fouled) and FE (least fouled). Earlier reports have further shown that the epibacterial community of *Fucus* spp. differ qualitatively and quantitatively among species in different levels of the intertidal zone (Lachnit et al., 2011; Stratil et al., 2013; Mensch et al., 2016; Quigley et al., 2020). This led us to hypothesize that the associated surface microbiota and metabolome may be the potential drivers of differential fouling intensity on the surfaces of three co-occuring *Fucus* spp. We first employed an untargeted metabolomics approach using the Feature Based Molecular Networking (FBMN) (Nothias et al., 2020) tool to comparatively analyze the algal surface metabolomes. We also comparatively characterized prokaryotic and eukaryotic communities associated with the thallus surface of the *Fucus* spp. by amplicon sequencing. Multivariate analyses, including linear discriminant analysis, allowed to infer significant markers driving the chemical and microbial differences on the surfaces of the *Fucus* spp.

## 2. Materials and methods

### 2.1. Sampling and sample processing

Fresh seaweed materials were sampled from Kiel Fjord, in the vicinity of the Bülk lighthouse (54°27′15.6′N and 10°11′55.0′E, Figure S1) in July 2019. *Fucus vesiculosus* (FV) was collected from max. 0.5 m depth, whereas *F. distichus* subsp. *evanescens* (FE) and *F. serratus* (FS) were collected at a maximum depth of 1 m from the water surface. Air and water temperatures were 16 °C and 17.5 °C, respectively, and water salinity was 13.1 PSU at a pH of 7. Following collection, algal materials were placed separately into sterile plastic bags. Ambient seawater (SW) was collected into sterile 1L Schott bottles. The biofilm on the surface of a stone (BF) close to the algae was also sampled into sterile 50 mL Falcon tubes. All samples were transported to the laboratory in a cooling box and processed on the same day. Algal samples were carefully rinsed with artificial seawater prepared by dissolving 1.8% Instant Ocean® in milliQ water (Arium® Lab water systems, Sartorius) to remove surface debris and loosely attached macrofoulers.

For analysis of the epiphytic microbial community, thallus surfaces of each *Fucus* sp. were sampled for amplicon sequencing of the conserved V3-V4 region of the bacterial 16S rRNA gene and the fungal internal transcribed spacer region. To avoid any change in the microbial community, samples for amplicon sequencing were processed immediately after arrival at the home laboratory. Surfaces of thalli were swabbed using a sterile cotton-tip and kept at -80 °C in sterile Eppendorf tubes. All algal samples were taken in triplicate, resulting in a total of 54 algal samples (18 samples per species). Ambient seawater (500 mL, SW), was filtered through cellulose nitrate filters (pore size 0.45µm, Whatman) in 6 replicates. Biofilm samples (Bf) were taken from a stone laying in the *Fucus* meadow in 6 replicates. All samples were stored at -80 °C prior to DNA extraction. Genomic DNA was extracted using the DNeasy PowerSoil Kit (Qiagen, Hilden, Germany). DNA concentration was measured prior to sequencing using a Nanovue UV-Vis spectrophotometer (GE Healthcare, USA) and stored at -80 °C.

### 2.2. Algal extractions

Algal surface extractions were carried out by both solvent dipping and solid-phase adsorption methods as described (Parrot et al., 2019)). Briefly, solvent dipping method was achieved by dipping the algal thalli into a stirring mixture of *n*-hexane:MeOH mixture (1:1) for 4 sec (1 kg algal material in 1 L solvent mixture). Crude extracts were filtered, *vacuum* dried and transferred onto a flash column packed with 30g of C18 material for desalting. After washing with 600 mL milliQ water^®^, it was eluted with 1200 mL of MeOH, *vacuum* dried to afford solvent dipping extracts (SD). For solid-phase (C-18) adsorption (SA), algal materials were covered with C18 material (Sepra C18-E, 50 µm, 65A, Phenomenex®, Torrance, CA, USA) and agitated (alga to C18 material ratio, 5:1). The C18 material was then washed into a glass column and sequentially eluted with 3-column-volumes of seawater, milliQ water® and 6-column-volumes of MeOH. The MeOH phase was dried under *vacuum* to yield solid-phase adsorption extracts (SA).

The algal material, which remained after surface extractions, and whole algal thalli (without surface extraction) were freeze-dried and pulverized by a speed rotor mill Pulverisette 14 (1.0 mm sieve ring, Fritsch GmbH, Idar-Oberstein, Germany). For each replicate (5 replicates per *Fucus* sp.), 4 g algal material was extracted by the Accelerated Solvent Extractor system ASE 350^TM^ (Dionex, Thermo Fisher Scientific, Sunnyvale, CA, USA). Extractions were performed by using MeOH and DCM as described previously (Heavisides et al., 2018). A three-step extraction was performed with water pre-rinse 10 min static (3 cycles), MeOH 5 min static (1 cycle) and DCM 5 min static (1 cycle). Samples were packed with acid-washed sand (Grüssing GmbH, Filsum, Germany) into 100 mL stainless steel cells and held at a temperature of 40 °C with a purge time of 250 sec and rinse volume of 30% cell volume. MeOH and DCM extracts were pooled, *vacuum* dried and named as surface-free extracts after C18 adsorption (SFA), surface-free extracts after solvent dipping (SFD) and whole algae extracts (W) without prior extraction of the surfaces.

### 2.3. Amplicon Sequencing and Bioinformatics

PCR, purification, library preparation and Illumina sequencing were implemented at Novogene Europe Ltd. (Cambridge, UK). Based on the unique barcodes, paired-end reads were assigned to samples before cutting off the barcodes and primer sequences. Paired-end reads were merged using FLASH (Version 1.2.11)(Magoč and Salzberg, 2011) to obtain Raw Tags which were filtered to obtain clean tags with fastp (Version 0.20.0). Cleaned sequences were further processed using the DADA2 package (version 1.26.0, (Callahan et al., 2016) in R. After filtering and trimming steps, forward and reverse reads were merged and chimeras were removed using the removeBimera function. For taxonomy assignment, the IdTaxa function in the DECIPHER package (version 2.26.0, (Murali et al., 2018) was used with SILVA_SSU_r138_2019 as reference database for bacterial sequences and UNITE_v2021_May2021 as reference database for ITS sequences. From the bacterial dataset, chloroplast sequences were removed (no mitochondrial sequences detected), and from the ITS dataset, sequences assigned to Anthophyta and Phaeophyceae were removed. Subsequent analysis was performed using the phyloseq package (v.1.42.0, (McMurdie and Holmes, 2013). Removal of samples below 15.000 reads and rarefying to 15.112 reads led to exclusion of two samples (one each from FS and FE) for further analysis of the 16S data. Due to low DNA concentrations, the ITS fragment was not sequenced for 4 samples from the BF and data reduction steps eliminated further two samples (one each from SW and FE) resulting in a total of 60 samples. ITS data were rarefied to 84.856 reads. For both datasets, alpha diversity was calculated for Observed ASVs and the Shannon index and statistically assessed using the Wilcoxon rank test. After determining the Bray Curtis dissimilarity index as best method to assess beta diversity, it was visualized by NMDS plots. Effects on community dissimilarity were analysed using permutational analysis of variance (PERMANOVA; Anderson, 2001) with the adonis2 function (vegan R package). Linear discriminant analysis Effect Size (LEfSe) was performed using microeco package (v.0.13.0, (Liu et al., 2021) and rarefaction curves were calculated using the microdev extension mecodev (v.2.0).

### 2.4. UPLC-QToF -MS/MS metabolomics

Aliquots (1 mg/mL in MeOH) of all extracts, W, SFA, SFD, SD, SA including solvent controls were injected (0.3 µL) into an Acquity UPLC I-Class system connected to a Xevo G2-XS QToF-MS (Waters®, Milford, MA, USA). A binary mobile phase (MP) comprised of ULC/MS grade solvents (VWR®, Pennsylvania, USA), water (A) and acetonitrile (B), both spiked with 0.1% formic acid (v/v)). Metabolite separation was achieved on a C18 column (Acquity UPLC HSS T3, 1.8 μm, 2.1×100mm, Waters®) at 40°C with MP infused at a flow rate of 0.6 mL/min and a gradient as follows: 1% B for 0.7min, increased to 30% B from 0.7 to 1 min, increased further to 99% B from 1 to 13.50, followed by an isocratic step of 99% B for 5 min, back to the initial conditions within 0.5 min and a column reconditioning step for 2 min for a total run time of 21 mins. MS was done in fast DDA acquisition mode with an ESI source over a mass range of *m/z* 50–1200 Da in both positive and negative polarities with a capillary voltage of 800 V, cone gas flow of 50 L/h, desolvation gas flow of 1000 L/h, source temperature of 150 ◦C, desolvation temperature of 500◦C with sampling cone and source offset at 40 and 80, respectively. The MS/MS experiment was achieved by using a collision energy ramp with the following settings: LM CE ramp start = 20, LM CE ramp End = 40, HM CE ramp start = 60, HM CE ramp End = 80 and a scan rate of 0.1 sec in centroid data format. All measurements were performed in quadruplicate and mass-corrected with LockSpray (Reference masses: 120.0813 and 556.2771 Da MSMS for Leucine enkephalin). Data from the positive polarity mode were more diverse and so were used for statistics. All data were analyzed using MassLynx® software (v4.2).

### 2.5. Molecular networking

ProteoWizard msconvert (version 3.0.20033; Vanderbilt University, Nashville, TN, USA) (Chambers et al., 2012) was employed to convert all .raw data files from UPLC-MS/MS to mzXML format and then imported and processed in MZmine 2.33 (Pluskal et al., 2010). The metabolomic features including retention time, *m/z* and peak areas were exported as .csv (feature quantitative table) while MS^2^ data were exported as .mgf files. The pre-processed data (.csv and .mgf files) were uploaded onto the GNPS platform (Wang et al., 2016) to generate molecular networks using the Feature-Based Molecular Networking (FBMN) workflow (Nothias et al., 2020). Here, identical MS/MS spectra are combined into ‘consensus’ spectra and displayed as nodes. The nodes are then linked with edges based on the similarity between the consensus spectra by spectral alignment algorithms. A molecular network was created with precursor ion and fragment ion mass tolerances set at 0.05 Da with edges filtered to have a cosine score above 0.7. Spectra were also searched against GNPS spectral libraries (score > 0.6) (Horai et al., 2010). The spectral network was further subjected to in silico tools; network annotation propagation (NAP) (da Silva et al., 2018) and Dereplicator+ (Mohimani et al., 2018), to enhance the results of the annotations. The outputs from molecular networking, NAP and dereplicator+ were merged through the MolNetEnhancer (Ernst et al., 2019) workflow which employs ClassyFire (Feunang et al., 2016) chemical taxonomy for chemical class annotation. The integrated network was visualized and analyzed by Cytoscape version 3.7.2 (Shannon et al., 2003). Additionally, peak ions were manually analyzed using MassLynx version 4.2 to predict the molecular formulae and searched against databases; NP Atlas (https://www.npatlas.org (accessed on 3 March 2023)) and Dictionary of Natural Product (http://dnp.chemnetbase.com (accessed on 3 March 2023)).

### 2.6. Statistical Analysis

Multivariate statistical analysis of the metabolomics data (LC-(+)-ESI-MS) was achieved on MetaboAnalyst platform (version 5.0) (Pang et al., 2021). Sample data was normalized by median and scaled using Auto scaling (mean-centered and divided by the standard deviation of each variable) prior to statistical analysis and visualization. The PLS-DA model generates variable importance in projection (VIP) scores which was used for biomarker prediction (Paix et al., 2019).

## 3. Results

### 3.1. Macroscopic fouling intensity

All three *Fucus* spp., i.e., *F. vesiculosus* (FV), *F. serratus* (FS) and *F. distichus* subsp. *evanescens* (FE) were sourced from the same location in the Kiel Fjord, Baltic Sea. FV was collected from max. 0.5 m depth, while FE and FS from max. 1 m water depth. As shown in Figure 1, different fouling intensity was observed visually on different species; the thalli of FV were the most intensely fouled, FS appeared to be relatively less fouled and FE was the cleanest (least fouled).

**Figure 1:**
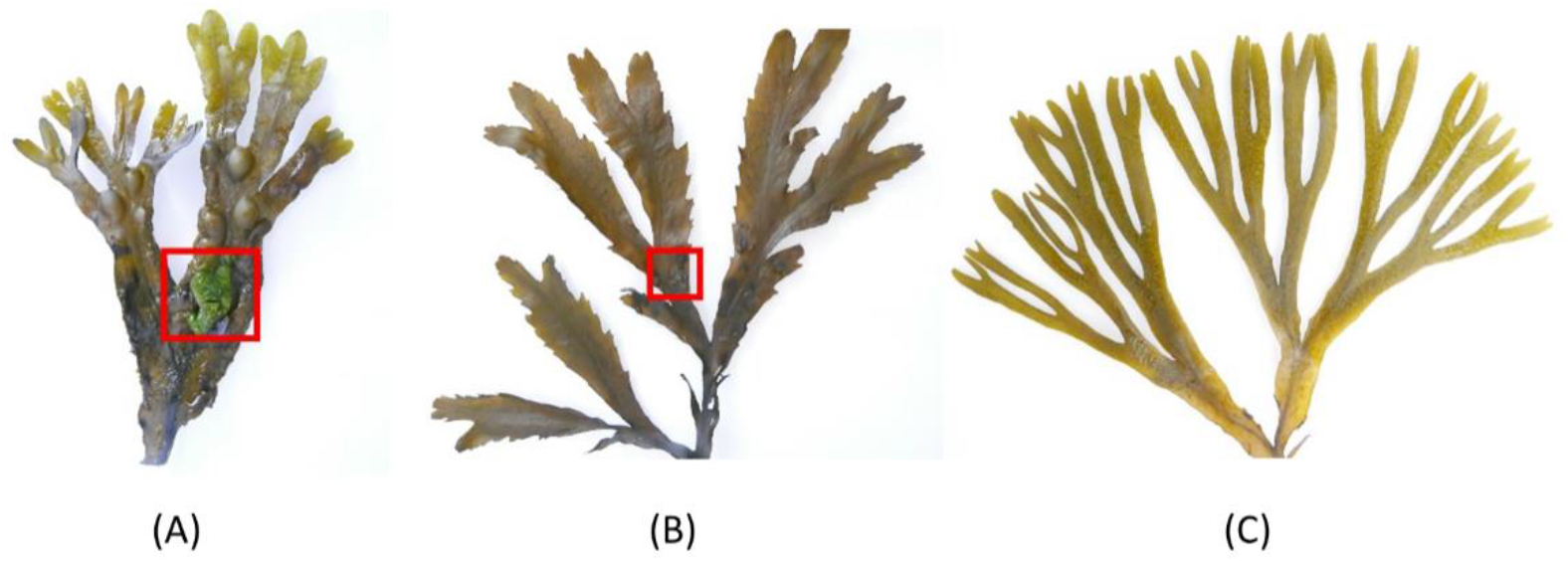
Macroscopic images of the thalli of three *Fucus* species sampled at Kiel Fjord. (A) The most intensely fouled *F. vesiculosus* (FV) incl. macrofouling as highlighted by the red square, (B) less fouled *F. serratus* (FS) with a small fouled area marked by the red square and (C) the least fouled *F. distichus* subsp. *evanescens* (FE)

### 3.2. Algal extractions

In order to analyze the whole surface metabolomes of the *Fucus* spp., two common surface extraction methods were employed, i.e., dipping in organic solvents and adsorption to a solid phase. Solvent dipping (SD) involved a very brief (4 sec) immersion of the algal thalli in a 1:1(v/v) *n*-hexane:MeOH mixture without leaching of compounds from the algal inner tissue (Rickert et al., 2016b; Parrot et al., 2019). The solid-phase adsorption (SA) consisted of mixing of the algal thalli with C-18 material to adsorb surface metabolites onto the C-18 followed by elution with MeOH (Cirri et al., 2016; Parrot et al., 2019). In parallel, the whole- and surface-free algal extracts were obtained to infer the biological origin of the metabolites observed on the algal surfaces. For this aim, a highly reproducible technique with automated Accelerated Solvent Extraction (ASE) was employed to obtain extracts of the whole untreated algae (W) and surface free algae after solvent dipping (SFD) and C18-adsorption (SFA).

### 3.3. Comparative untargeted metabolomics of crude extracts

All extracts were analyzed by ultra-high-resolution mass spectrometry (UPLC-MS) in both positive (+) and negative (-) ion modes. Manual inspection of the acquired UPLC-MS chromatograms revealed more complex profiles in the positive ion mode than in the negative ion mode. The acquired UPLC-MS (+) chromatograms revealed fewer peaks in the C18 SA extracts (Figure S1) compared to the SD extracts (Figure S2) of all three *Fucus* spp. Also, metabolite compositions of the surface free extracts (SFA: Figure S3, and SFD: Figure S4) were similar to the whole algal extracts (W, Figure S5). MZmine pre-processing of all raw MS (+) data resulted in 6,325 *m/z* features. After applying several filtering steps (incl. removal of solvent-derived features), the final data set comprised 366 features from all extracts (Table S1).

#### 3.3.1. Comparison of the surface metabolomes of SA and SD

Surface extracts represented the focus of analyses, as their constituents are considered to have the largest impact on epibiosis and fouling on the algal surface. Two surface extraction protocols revealed a vast difference in the number of observed metabolites. SD method produced 268 features while SA generated 39 features (Figure 2A). SA shared 90% of its features (35) with SD and had only 4 unique features, putatively annotated to 4-pyridoxic acid, a sphingolipid and two unannotated ions (Table S1). SD extracts contained 233 unique features (Figure 2A). The feature distribution among species, based on surface extraction techniques, are displayed in the Figures 2B and 2C. FS showed the highest number of features in both extraction methods.

**Figure 2:**
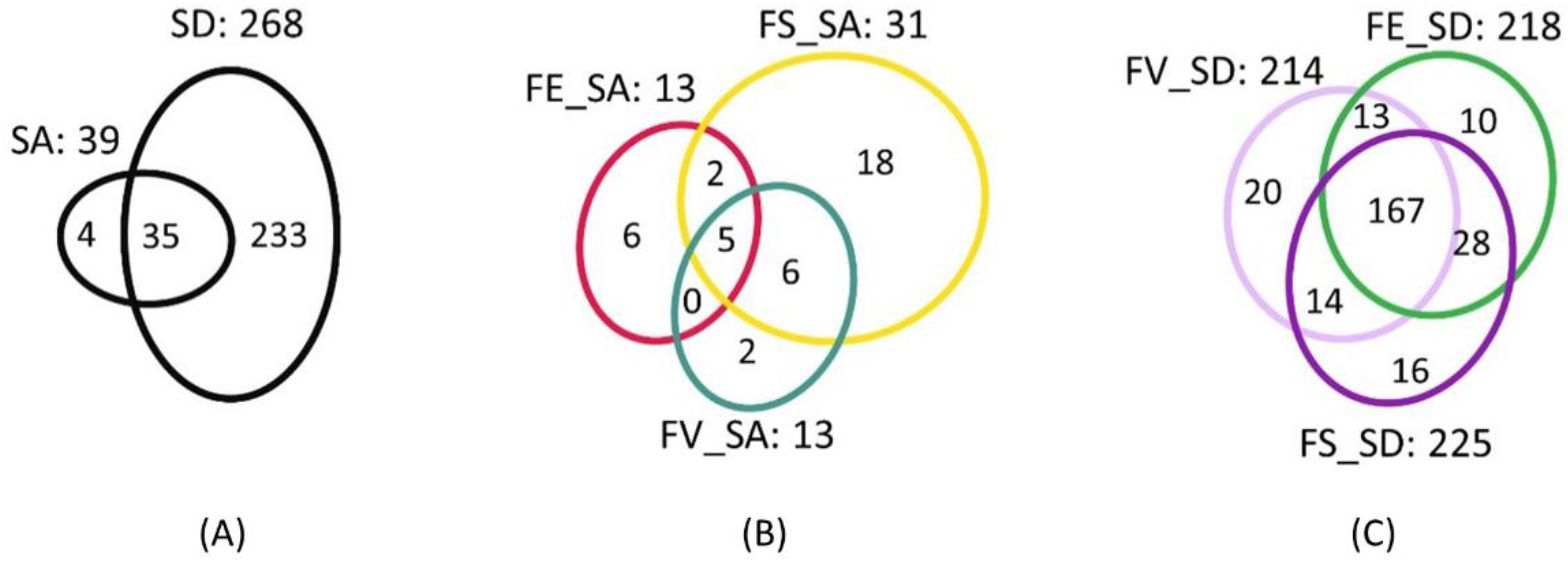
Venn diagrams displaying the distribution of features between **(A)** surface adsorption (SA) and solvent dipping (SD) methods **(B)** *F. vesiculosus* (FV), *F. serratus* (FS) and *F. evanescens* (FE) by SA and **(C)** FV, FS and FE by SD.

In order to facilitate annotation of significant features and the surface metabolomes, Feature-Based Molecular Networking (FBMN) via the GNPS platform (Wang et al., 2016) was performed to organize the UPLC-(+)-MS/MS data of extracts into clustered ions of similar MS/MS spectra. An extra network using data obtained in the negative ion mode was also generated (Figure S6). Only annotations manually confirmed by the molecular formula prediction using MassLynx®, fragment verification and the source of the hit compound are displayed in the MNs.

The MN of the surface metabolomes comprised of 272 nodes in 24 clusters (Figure 3A). Node size, a reflection of the sum of precursor intensity, allowed for visualization of the most abundant compounds within clusters, while the pie chart in node shows the relative abundance of the metabolite in each *Fucus* sp. The largest cluster of the surface metabolome displayed defined product ions either at *m/z* 212.2401 or *m/z* 240.2682 in all 3 spp. However, manual dereplication did not provide any structural information and hence annotation to individual ions was not possible. MolNetEnhancer with *in silico* tool NAP annotated the cluster as ‘carboxylic acid and derivative’ class of compounds (Figure 3A and Table S1). Polar lipids such as betaine lipids and galactolipids were putatively identified as dominant metabolites across the surfaces of the *Fucus* spp. These included the betaine lipids diacylglyceryl trimethyl-homoserine (DGTS), monoacylglyceryl hydroxymethyl-trimethyl-β-alanine (MGTA), and galactolipids monogalactosyl diacylglycerol (MGDG), digalactosyl diacylglycerol (DGDG) (Figure 3A and Table S1). Also, the tetrapyrrole class of compounds was shared by all three *Fucus* spp. These clusters are chlorophyll-related and prevalent in microalgae residing on the seaweed surfaces (da Silva and Lombardi, 2020). Other metabolites putatively identified on the surfaces include the carotenoids; fucoxanthinol, dehydrated fucoxanthin and other cluster-members, shared by all species. Some singletons were annotated; the sugar-polyol mannitol, amino acid *L*-tryptophan, and the betaine type amino acid ulvaline, were also shared by all seaweed species, albeit at different concentrations. In the negative ion mode, we could annotate 4 nodes in the sulfolipids cluster (SQDG), 2 nodes in the phosphatidylethanoloamine cluster and a node in the lyso phosphatidylinositol cluster (Figure S6), shared by the *Fucus* spp.

**Figure 3:**
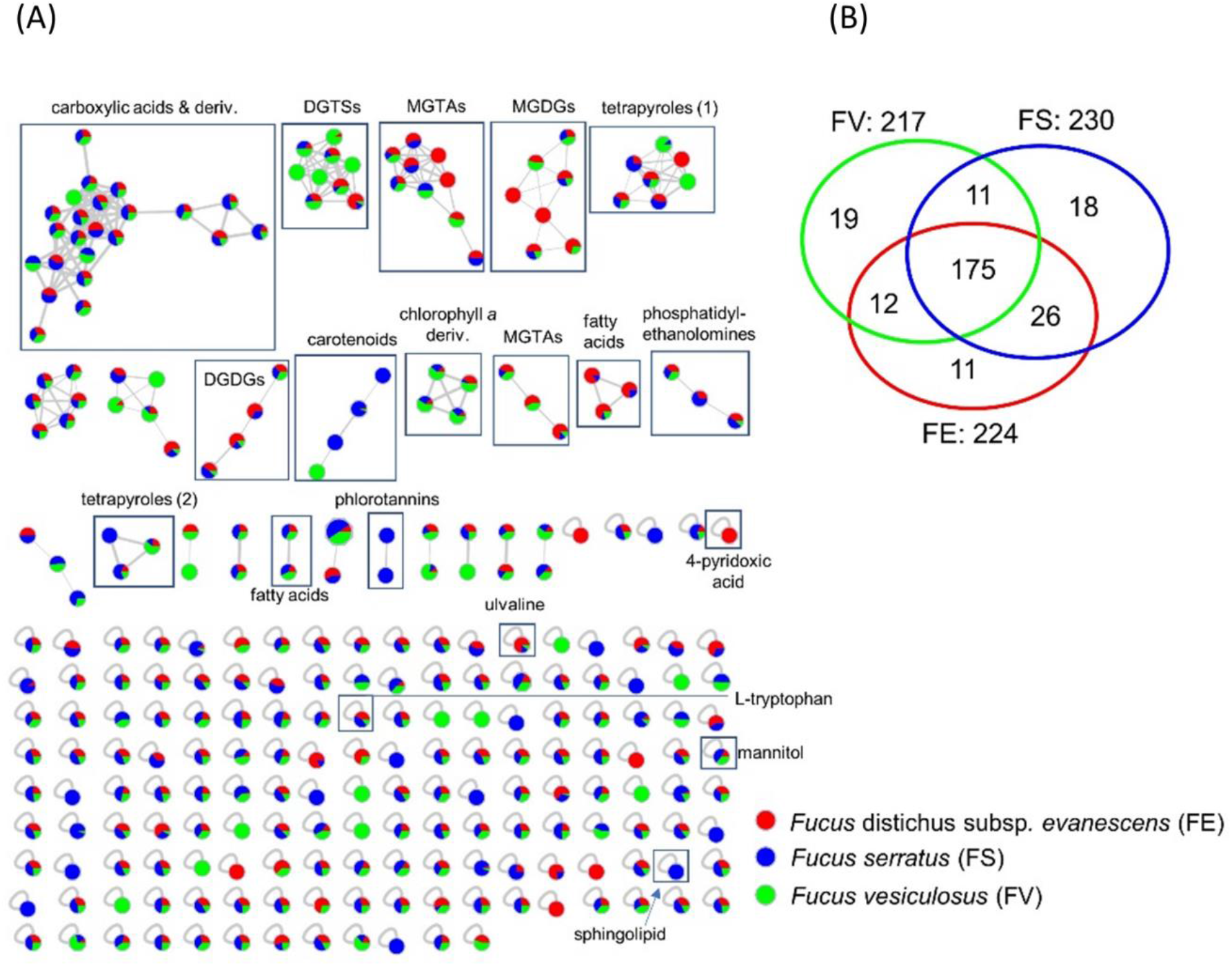
(A) MN generated from UPLC-(+)-ESI-MS/MS data of all algal surface extracts. Node sizes are modulated according to the sum of intensities of the ions in all extracts while the colors in the pie chart of each node represents the relative quantity of the ion from FE (red), FS (blue) and FV (green). (B) Venn diagram displaying the nodes distribution among species from the combined surface extracts. Betaine lipids diacylglyceryl trimethyl-homoserine (DGTS), monoacylglyceryl hydroxymethyl-trimethyl-β-alanine (MGTA). Galactolipids monogalactosyl diacylglycerol (MGDG), digalactosyl diacylglycerol (DGDG)

As shown in Figure 3B, SA and SD extracts of FS showed the highest number of metabolites (230 nodes), while FE and FV recorded 224 and 217 nodes, respectively. A high proportion of nodes (80%) were shared, while a total of 48 nodes were unique to the surfaces of each *Fucus* sp. (18 nodes for FS, 11 for FE and 19 for FV (Figure 3B). Of these, 6 metabolites exclusive to the surface of FE were annotated as betaine lipids MGTA 20:5, MGTA 18:3, galactolipid MGDG 36:4, sugar-polyol 1-O-β-D-glucopyranosyl-D-mannitol, pyridine 4-pyridoxic acid and the fatty acid eicosatetraenoic acid derivative. FV also showed exclusive betaine lipids including DGTS 32:3, DGTS 32:2 and DGTS 34:4, galactolipid MGDG (18:4) and the carotenoid fucoxanthin dehydrated (5 annotations). Eight compounds exclusive to FS surface were annotated; sphingolipids N-(1,3-dihydroxyoctadecan-2-yl)acetamide and C20-phytosphingosine, carotenoid fucoxanthinol, tetrapyrrole phaeophorbide A methyl ester and three phlorotannins fucodiphloroethol (A and/or B) and fucodiphlorethol (A or B).

Next, we performed multivariate analysis, i.e., supervised partial least squares discriminatory analysis (PLS-DA) in order to determine the variations and identify potential marker compounds in the surface metabolomes. PLS-DA revealed a clear discrimination among the surface extracts indicating a total variance of 28.2% (Figure 4A, [Permutation test result; p<0.05]). A hierarchical clustering heatmap was performed to visualize different concentrations of the metabolites on the surfaces of the *Fucus* spp. (Figure 4B). Several regions highlighted in green, blue and orange represent metabolites in higher abundances relative to the other regions. The most important metabolites responsible for the algal surface discrimination are highlighted by variable importance in projection (VIP) scores (Figure 4C). These were betaine lipids; MGTA 20:4, MGTA 18:1, DGTSA and ulvaline, most abundant on the surface of FE, while the galactolipid MGDG 34:4 was abundant on FV surface. The carboxylic acid derivative (C.A. deriv.) class of compounds were also significant on FV (SD) and FE (SA) (Figure 4C and Figure S7).

**Figure 4:**
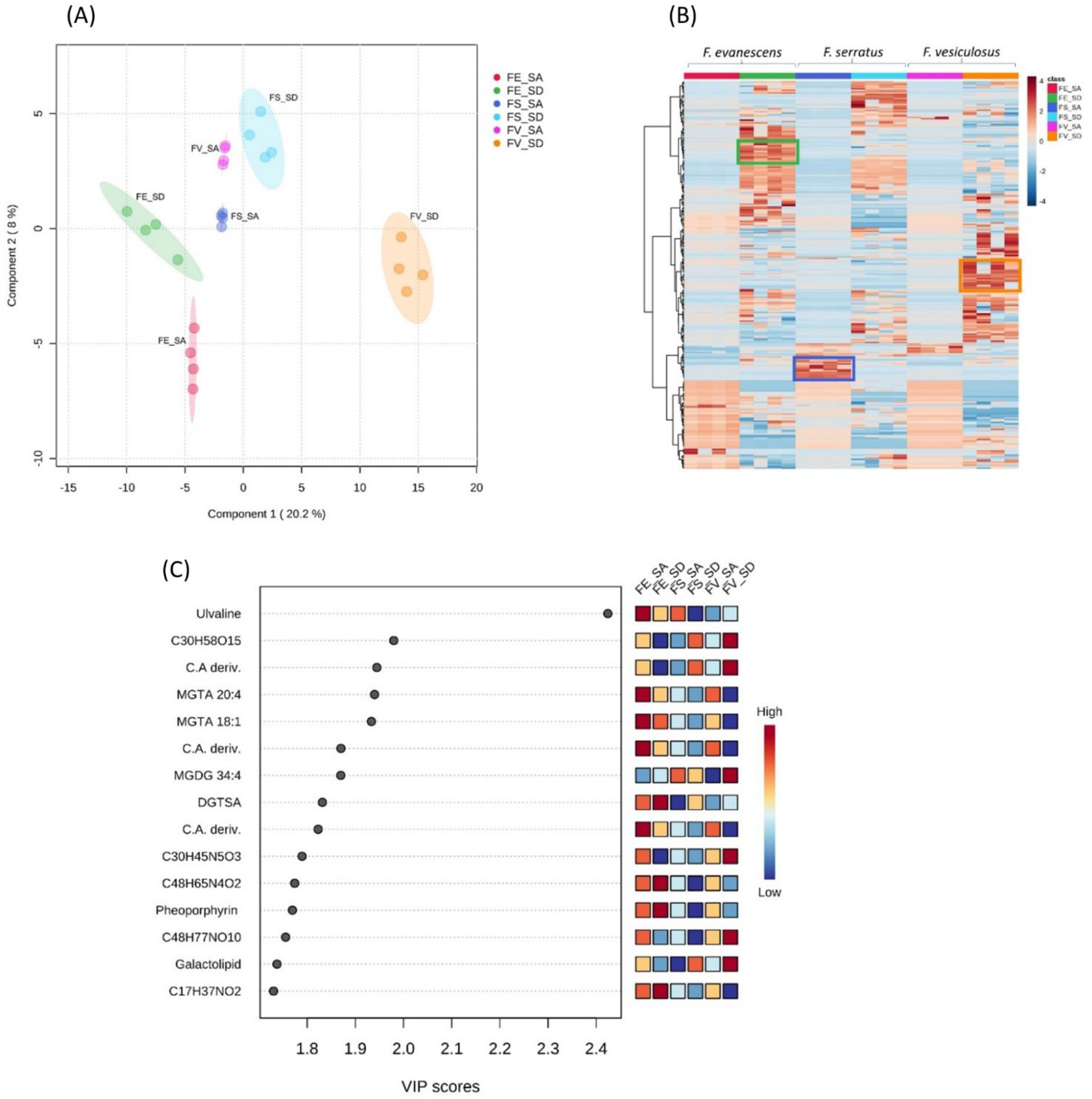
Multivariate analyses of Fucus spp. (A) PLS-DA scores plot generated from UPLC-(+)-ESI-MS data of all surface extracts of *Fucus* spp. obtained from C18-adsorption (SA) and solvent dipping (SD) methods. (B) Heatmap visualization of the different concentrations of the metabolites, obtained from UPLC-(+)-ESI-MS data, expressed on the surfaces of the three *Fucus* spp. Columns: samples; Rows: metabolites; Color indicates metabolite expression values, blue: lowest, red: highest. SA and SD extracts for *F.* distichus subsp. *evanescens* (FE_SA/FE_SD), *F. vesiculosus* (FV_SA/FV_SD) and *F. serratus* (FS_SA/ FS_SD) Highlighted regions green, blue and orange represents metabolites in higher abundances (C) Top 15 annotated metabolites ranked by VIP values. The mini heatmap on the right indicates their concentrations on the surface extracts of the *Fucus* spp.

Comparing the surface extracts (SA and SD) to the whole (W) and surface-free extracts (SFA and SFD), we observed a clear separation/difference between the surface metabolites (SA, SD) and the algal inner tissue metabolites (W, SFA, SFD) (Figure S8). Based on the MN (Figure S9) the main difference between the surface metabolomes (and inner tissue metabolomes) was the unannotated ‘carboxylic acid and derivatives’ cluster, which was only detected in surface extracts indicating its origin from the surfaces and shared by all *Fucus* spp. All annotations made are displayed in Table S1.

### 3.4. Microbiome analysis

The surface microbiomes of the three *Fucus* spp. were comparatively analyzed by amplicon sequencing of the V3-V4 hypervariable region of the prokaryotic 16S rRNA gene and the eukaryotic internal transcribed spacer 2 region (ITS-2), in order to determine if the epiphytic communities were species-specific and could illuminate differential fouling levels. Seawater (SW) and biofilm from a stone (BF) taken from within the *Fucus* meadow were included as references for comparisons.

#### 3.4.1. Epiphytic bacterial microbiome

The Illumina NovaSeq sequencing platform generated a total of 7,193,202 raw paired-end reads out of 66 samples (*Fucus* surface samples and controls incl. replicates) in total. These were reduced to 2,096.951 bacterial reads after quality filtering, chimera and chloroplast sequence removal. Rarefaction analysis indicated sufficient sequencing depth (Figure S10). The highest alpha diversity of all algal species is observed in FV, followed by FE and the lowest alpha diversity was in FS (Figure S11). Significantly distinct bacterial alpha diversity was observed (Wilcoxon rank sum test) between the pairs of FV and FS, and between FE and FS, but not between FV and FE and between reference samples SW and BF. Moreover, highly significant variations in alpha diversity were observed for FS and FE to BF and SW, whereas FV showed only slight (to BF) or no (to SW) significant differences. No significant alterations were observed between different individuals (Figure S12).

In total, 15 bacterial phyla (with abundance > 1%) were observed in the samples (Figure 5A). Epibionts of all three *Fucus* spp. and the reference samples were dominated by three bacterial phyla, Proteobacteria, Cyanobacteria and Actinobacteria, which constituted 89.8 – 96.2% of the community in all samples. The highest abundance of Proteobacteria was in SW (67.3%) and the lowest on surface of FS (42.1%). The highest Cyanobacteria abundance was detected in BF (34.5%) and the lowest on FV (18.2%). Actinobacteria were most abundant on FS (24.6%). Surprisingly, the phylum Bacteroidota was only detected as rather minor constituent of the bacterial community in all samples with highest abundance on FE (0.12%). The extremophilic bacterial phylum Deinococcota was specific for seaweeds, with highest abundance on FS (4.2%). The dominant genus on the surfaces of the seaweeds was *Schizothrix* (Cyanobacteria); 24.41% on FV, 29.61% on FS, and 26.02% on FE (Table S2).

**Figure 5:**
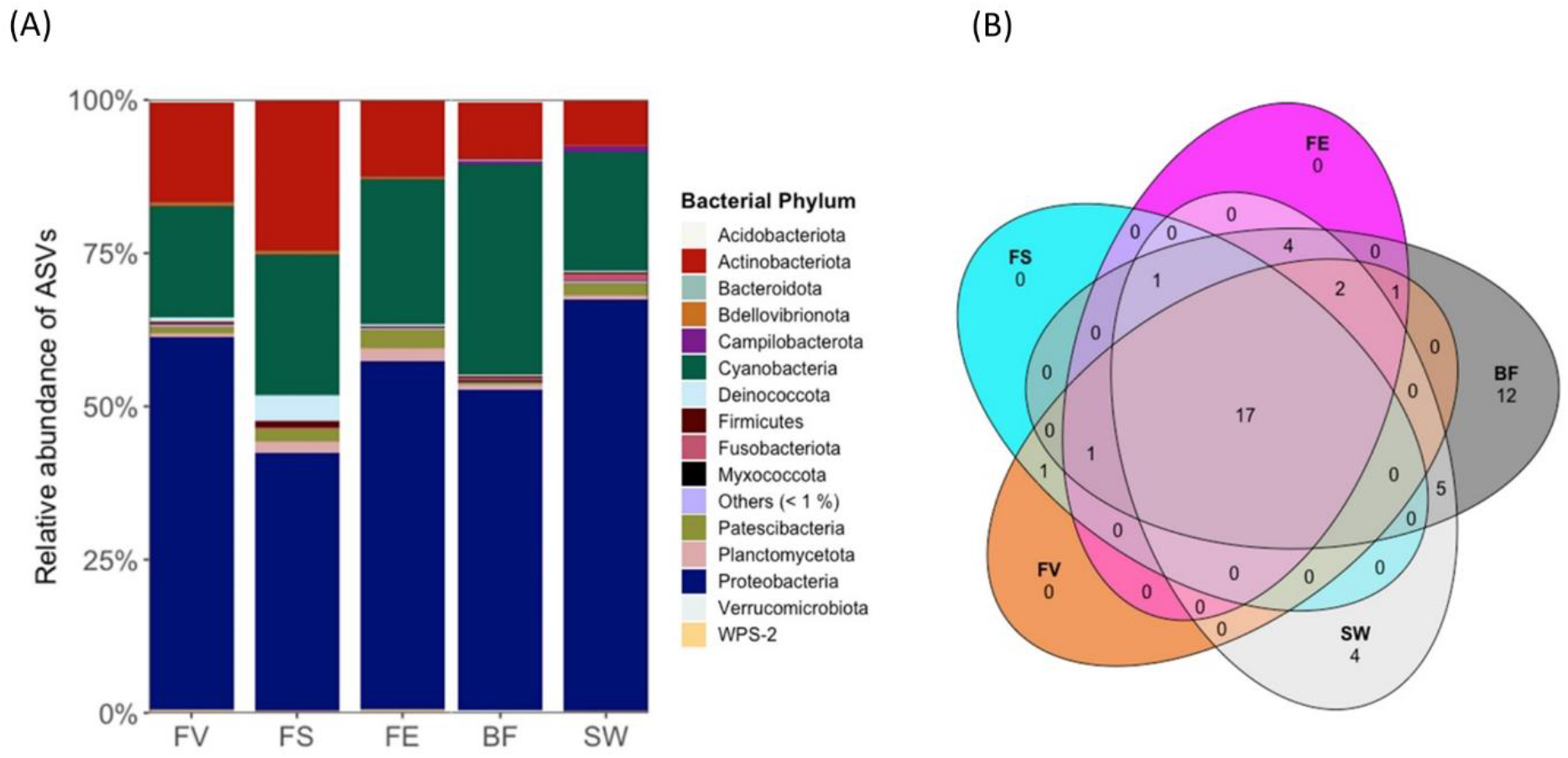
(A) Bacterial phyla associated to surfaces of *Fucus* spp. and stone biofilm (BF) and seawater (SW) references. FV: *Fucus vesiculosus*, FS: *F. serratus,* FE*: F. distichus* subsp. *evanescens.* Others (<1%) represents several phyla with less than 1% relative abundance. (B) Venn diagram of bacterial amplicon sequences at genus level.

Reference SW and BF showed bacterial community differences from the algal surfaces. The phyla Acidobacteriota and Firmicutes were exclusive to BF and SW. More so, *Clade Ia* (Proteobacteria) and *Phormidesmis* (Cyanobacteria) represented the most abundant genera in SW (22.46%) and BF (12.76%) respectively (Table S2). As shown in the Venn diagram (Figure 5B), 17 genera were identified in all samples, and no genus was specific to any of the algal surfaces. Beta diversity analysis using Bray-Curtis distances highlighted the dissimilarity of bacterial communities on each seaweed surface and reference samples (Figure S13, Table S3). The distribution of the bacterial order for the *Fucus* spp. is given in Figure S14.

In order to determine the taxa most likely responsible for the observed variations, a Linear discriminant analysis Effect Size (LEfSe) was performed using the LEfSe function implemented in R microeco package (v. 0.13.0) using a threshold value of 4.

This identified 3 biomarkers for FE, 7 for FS, and 6 for FV as taxa responsible for the differences in the three *Fucus* surface microbiome. Significantly abundant epiphytic bacteria (biomarkers) on FE were the genus *Robiginitomaculum* (Family: Hyphomonadaceae, Order: Caulobacterales, Phylum: Pseudomonadota) and genus *Yoonia-Loktanella* (Phylum: Pseudomonadota). The most significant discriminant genera on the surface of FS were *Truepera* (Family: Trueperaceae, Order: Trueperales, Phylum: Deinococcota), *Litorimonas* (Phylum: Pseudomonadota) and a member of the order Deinococcales (Class: Deinococci, Phylum: Deinococcota). Octadecabacter (Family: Roseobacteraceae, Order: Rhodobacterales, Phylum: Pseudomonadota), Microtrichaceae (Phylum: Actinobacteriota) and Sva0996 marine group (Phylum: Actinomycetota) were the significant discriminant taxa on the surface of FV (Figure 6). For the controls, 12 and 16 biomarkers were identified in the BF and SW respectively (Figure 6).

**Figure 6:**
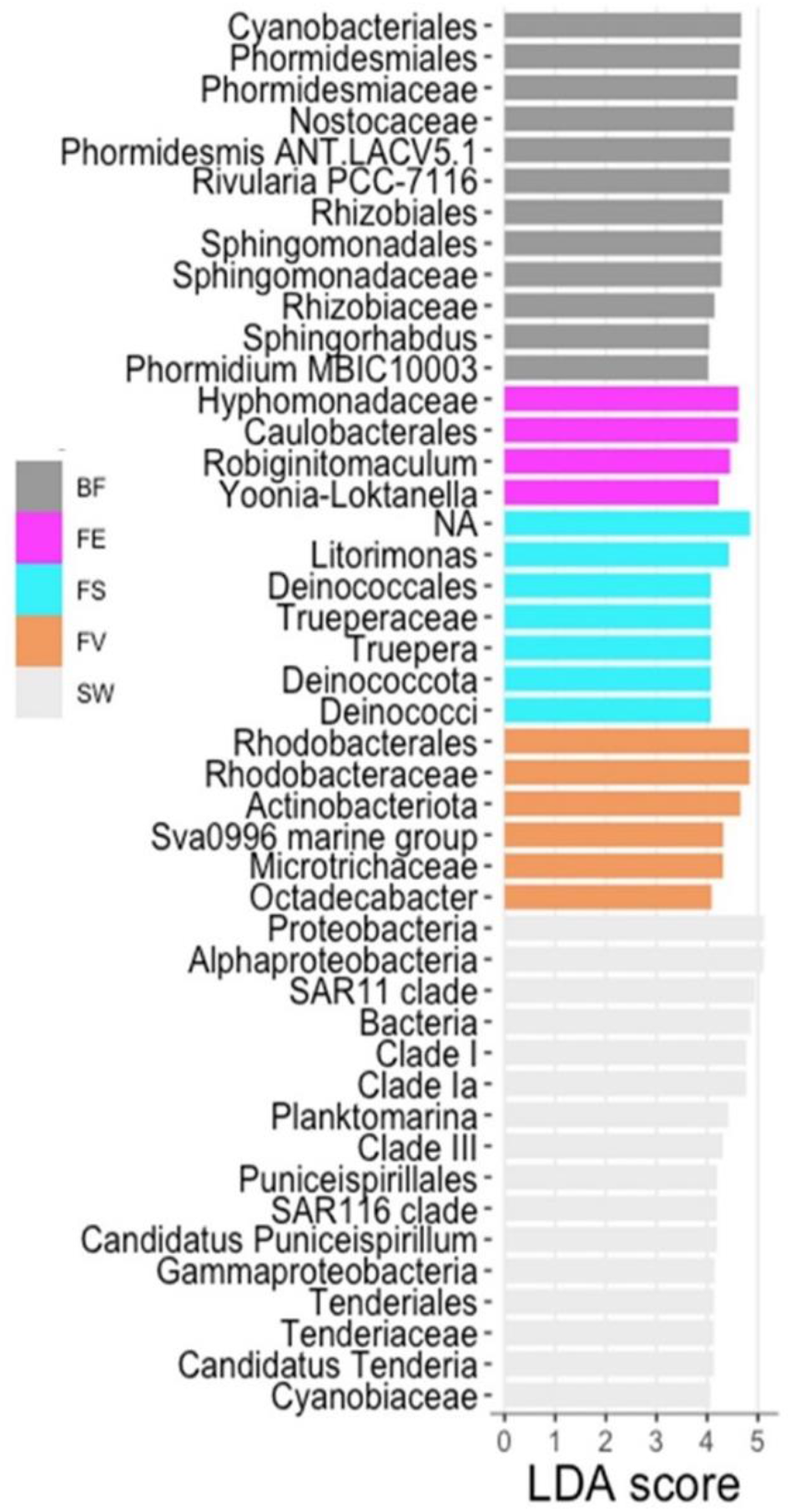
Bar plots showing linear discriminant analysis (LDA, threshold: 4) score after differential abundance analysis of bacterial taxa by LEfse according to origin, FV: *Fucus vesiculosus*, FS: *F. serratus,* FE*: F. distichus* subsp. *evanescens*, BF: biofilm on stone, SW: seawater.

#### 3.4.2. Comparative analysis of eukaryotic epiphytes

For the eukaryotic community, a total of 6.824.917 raw reads were obtained from 60 samples as 6 samples (4 BF, 1 SW and 1 FE samples) failed quality checks/filtering. Quality filtering and chimera removal resulted in 6.053.795 reads and 1962 taxa for phyloseq analysis. Rarefaction analysis also revealed sufficient sequencing depth (Figure S15). Analysis of alpha diversity using the Shannon index as well as the ASV diversity showed a considerably lower ITS sequence diversity on all seaweed surfaces compared to SW and BF (Figure S16). Slightly significant differences were observed between FV/FS and FV/FE but not between FS/FE and the references SW/BF. ASV diversity was insignificant between individuals (Figure S17).

The eukaryotic community was dominated by the phyla Ciliophora, Chlorophyta and Ascomycota on all algal surfaces (Figure 7). Cnidaria and Ascomycota were more abundant on FE and FS than on FV, and, the relative abundance of Mucoromycota on FE was higher than on FS and FV. Fungal phylum Ascomycota was by far the most abundant phylum in BF, followed by Chlorophyta. SW was largely dominated by the phyla Ciliophora and Cnidaria. The relative abundances of eukaryotes at the genus level are displayed in Table S4. Except BF, fungi accounted for 4-45% of the ITS sequences. A considerably low frequency of fungi was observed on FS. On FE, Ascomycota were most abundant followed by Mucoromycota. Basidiomycota were only detected on FV surfaces in notable abundances. PERMANOVA analysis of the whole dataset showed significant differences with regard to sample origin and individuals (Table S5).

**Figure 7:**
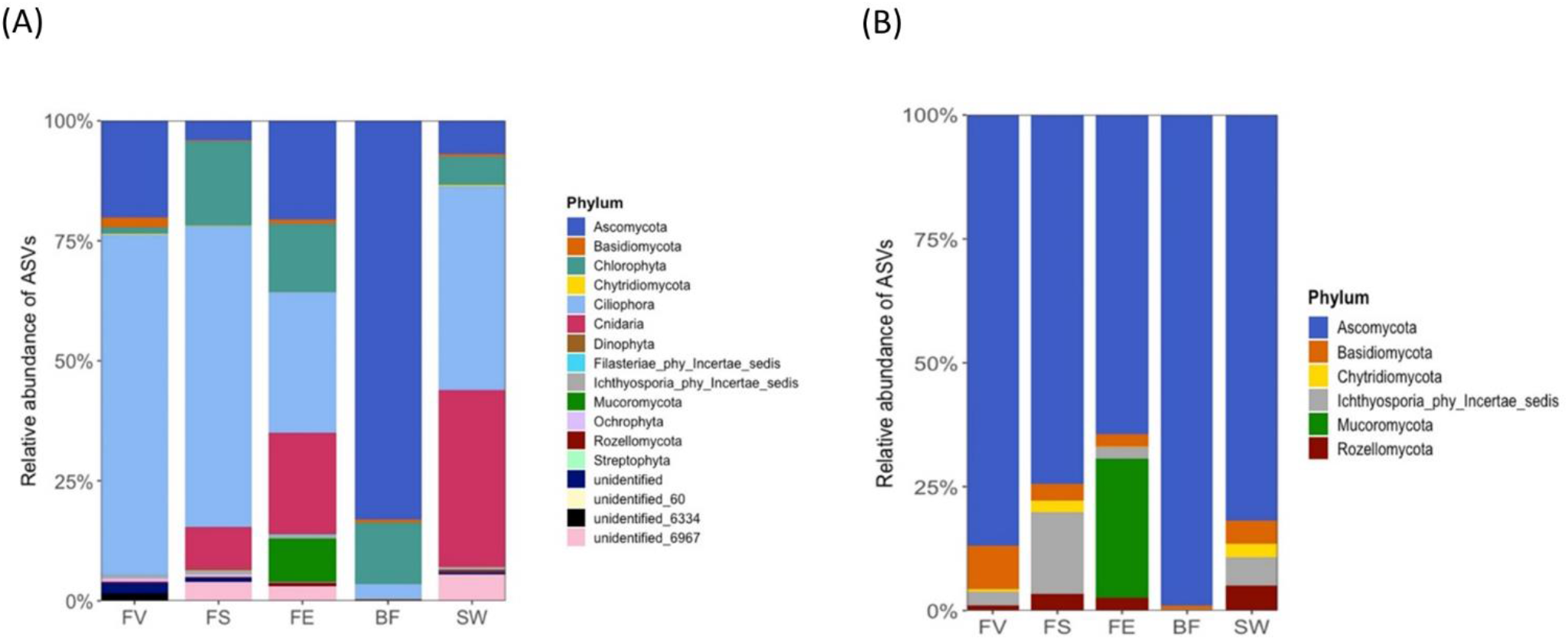
Community composition of epiphytic eukaryotes on *Fucus* spp. based on amplicon sequencing of the ITS region. (A) Eukaryotic community composition at phylum level. (B) Fungal community composition at phylum level. FV: *Fucus vesiculosus*, FS: *F. serratus,* FE*: F. distichus* subsp. *evanescens,* BF: biofilm on stone, SW: seawater.

With reference to fungal-derived sequences, alpha diversity analysis (ASVs) showed only significant differences between FV and FS and of all algae to SW and BF (Figures S18 and S19). Ascomycota, Basidiomycota, Ichthyosporia and Rozellomycota were the most abundant phyla on the algal surfaces (Figure 7B). FE surfaces exclusively harbored Mucoromycota. Differential abundances of fungi were observed at order and genus levels (Figure S20, Table S6).

The most dominant fungal genera on FV were *Candida* (82.30%), and *Haptocillium* (9.20%), while *Sphaeroforma* (38.04%) and *Alternaria* (25.00%) were most abundant on FS (Table S6). *Mucor* represented the most abundant genus on FE (65.41%) followed by *Alternaria* (14.91%) (Table S6). Due to the reduced sample set (12 samples removed, total of 189 taxa), beta diversity analysis and statistical assessment was not successful for the fungal subset. However, LEfSe analysis identified 10 biomarkers for BF, 6 for FE and 4 for FS. No differentially abundant taxa were identified for SW and FV (Figure 8). Fungi of the orders Sporidiobolales (Division: Basidiomycota) and Dothideales (Division: Ascomycota) were enriched on the surface of FE. On FS, Erysiphales (Class: Erysiphaceae, Division: Ascomycota), Eurotiomycetes (Division: Ascomycota) and fungi of the division Chydridiomycota were enriched (Figure 8).

**Figure 8:**
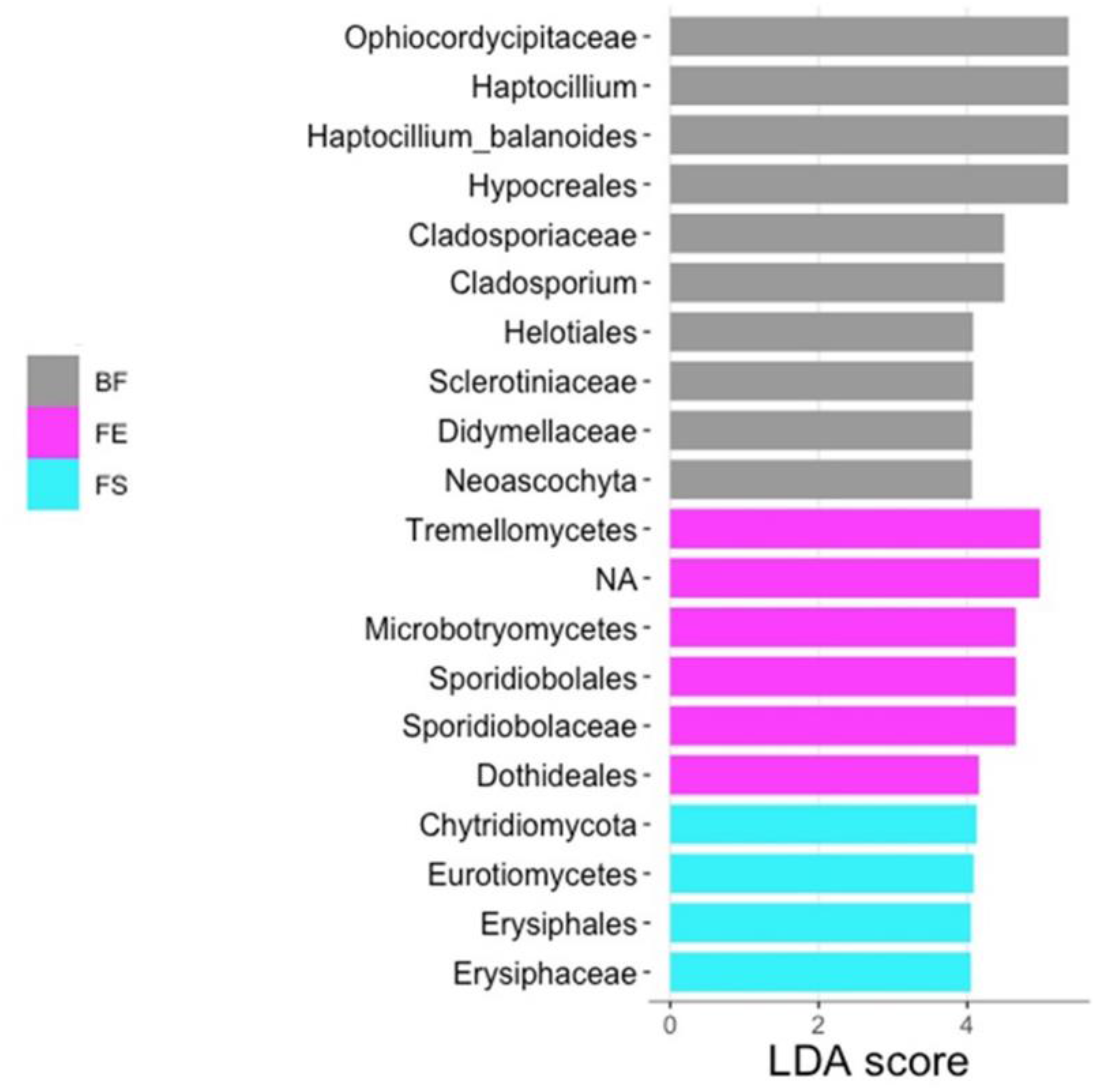
Differential abundance by Linear discriminant analysis (LDA, threshold: 4) of fungal taxa based on ITS amplicon sequences according to sample origin. FV: *Fucus vesiculosus*, FS: *F. serratus,* FE*: F. distichus* subsp. *evanescens,* BF: biofilm on stone, SW: seawater. No differentially abundant taxa were identified for SW and FV.

## 4. Discussion

There is ample evidence suggesting that seaweeds are able to control and shape their surface epibiome, which is considerably influenced by environmental gradients (Wahl et al., 2010; Lachnit et al., 2013; Pantos et al., 2015). It was intriguing to visually observe differential fouling on the thalli of three-foundation species, *F. distichus* subsp. *evanescens* (FE), *F. serratus* (FS) and *F. vesiculosus* (FV) co-occurring in the Kiel Fjord. We aimed to understand several factors that may drive different fouling levels on the seaweed surfaces. The composition of surface metabolome and microbial community are pivotal in biofilm formation/epibiosis (da Gama et al., 2014; Saha et al., 2018). Therefore, identification of the unique core taxa of the surface microbiome, and metabolome associated with the surfaces of the foundation species were of interest. Although the thalli surface metabolomes were the main target of this study, the whole biomass and surface-free algal materials were also extracted and analyzed as controls. The whole algal extracts were analyzed to index the general metabolic profile, while the surface free algal extracts served assessing the residual metabolome after removal of the surface metabolome. Many known *Fucus* derived compounds, with reported diverse biological activities, were annotated in all the three species (Table S1).

For the surface extracts, two extraction protocols previously described, were employed (Cirri et al., 2016; Rickert et al., 2016a; Parrot et al., 2019; Papazian et al., 2019). Although SA produced fewer compounds (14% of the total surface metabolites), SA extract-specific compounds were annotated including the sphingolipid *N*-(1,3-dihydroxyoctadecan-2-yl)acetamide (Pendyala et al., 2003) and pyridoxic acid, a ROS (reactive oxygen species) lowering compound (Wilson et al., 2017; Latorre et al., 2019). Similarly, the surface quorum sensing metabolite N-(3-oxooctanoyl) homoserine lactone was extracted by SA only (absent in SD extract) from the surface of FV in a previous study (Parrot et al., 2019). Although this compound was not identified in the current study, the exclusive presence of some compounds in SA supports the complementarity of the two surface extraction techniques.

Our efforts to compare the surface metabolome of the *Fucus* species revealed diverse compounds as visualized in molecular networks (MN). Based on automated and manual dereplication, about 70% of extracted compounds were annotated to the class level. Indeed, the surface metabolome of FV has been extensively studied with reported surface metabolites including citric acid, the amino acids (*L*-serine, *L*-threonine, *L*-asparagine and *L*-proline), simple sugars (mannitol, glycitol and ribitol) and dimethylsulphopropionate (Lachnit et al., 2010; Rickert et al., 2016b). In the current study, the amino acid *L*-tryptophan and the sugar mannitol were annotated with the later reportedly represent approx. 30% of the dry weight of *Fucus* spp. (Groisillier et al., 2013; Menshova et al., 2016). Mannitol is an osmoregulatory and energy-storage compound utilized in brown algae in ways such as to scavenge oxygen radicals and produce chemical defenses during stress (Dittami et al., 2011; Rickert et al., 2015). Its production showed a significant negative correlation to microfouling on FV (Rickert et al., 2016a). It also displayed bacteriostatic activity against multiple *Bacillus* spp. (Singh, 2014), hence contribute to shaping algal surface microbial community esp. in FE which showed a surface-specific mannitol linked to a β-glucopyranose residue (1-O-β-D-glucopyranosyl-D-mannitol) in our study.

Our results show species-specific and shared compounds in significantly different quantities on the *Fucus* spp. which drive the chemical differences on the surfaces. The betaine lipids and galactolipids (lyso-type lipids) MGTA 20:5, MGTA 18:3 and MGDG 36:4, as well as MGTA 20:4 and MGTA 18:1 were identified as discriminant metabolites mostly abundant or specific on FE surface. These polar lipids have been associated with diverse bioactivities including antifouling, antimicrobial and osmoprotectant (Heavisides et al., 2018; Buedenbender et al., 2020; Plouguerné et al., 2020; Alagawany et al., 2022). The betaine-type amino acid ulvaline, first isolated from the microalgae *Monostroma nitidum* (Abe and Kaneda, 1975), was observed in significantly higher quantity on FE (FE_SA) than the other *Fucus* spp. Unfortunately, very little is known of its biological function, but similar to other betaine lipids, it may be relevant to marine algal metabolism and antioxidant effect during biotic stress (Llewellyn et al., 2015; Sofy et al., 2020). The putative vitamin B12 derivative 4-pyridoxic acid, with antioxidant property, was also observed only on the surface of FE and has also been previously identified in some seaweeds as either an exogenous molecule from associated bacteria or self-made (Croft et al., 2005). The bioactivities of these FE surface metabolites align with the least fouled surface of FE observed.

On the surface of FS, we putatively identified three phlorotannins. Phlorotannins are hydrophilic in nature and form part of the cell wall structures of brown algae, constituting about 5-12% of dry weight of *Fucus* spp. (Okeke et al., 2021). Among the wide range of biological activities, phlorotannins defend the algae against grazers (Koivikko et al., 2005; Negara et al., 2021) and may contribute to the less fouled surface of FS. Additionally, the antioxidant fucoxanthinol (deacetylated derivative of fucoxanthin (Sachindra et al., 2007)) was observed only on the surface of FS.

Similar to previous work, dehydrated fucoxanthin was observed only on the surface of FV (Parrot et al., 2019). Unlike FE, the surface of FV showed more diacyl-betaine lipids such as DGTS 32:3, DGTS 32:2 and DGTS 34:4. ‘Acyl editing’, the diacylation and reacylation cycles in lipid metabolism (Hoffmann and Shachar-Hill, 2023), could influence their biological activities. However, this remains to be investigated. Other discriminant compounds belonging to the carboxylic acid cluster could not be annotated.

As for the microbiome analysis, the overall prevalence of Proteobacteria, Cyanobacteria and Actinobacteria in the seawater, stone biofilm and on the surfaces of *Fucus* spp. is in line with other studies as they are usually dominant in marine environments. Proteobacteria are ubiquitous in the marine environment due to their metabolic versatility and ability to enhance surface colonization and biofilm formation (Bonilla-Rosso et al., 2012; Selvarajan et al., 2018). Our observation of a wide spread of Cyanobacteria on the surfaces of *Fucus* spp., collected during the summer season, is consistent with previous findings that showed the abundance of Cyanobacteria on FV in summer, but not in winter (Lachnit et al., 2011). Actinobacteria and members of the phyla Planctomycetota, Verrucomicrobiota and Fusobacteriota, observed in relatively lesser abundances, have previously been reported on many seaweed surfaces (Lachnit et al., 2011; Bondoso et al., 2014; Selvarajan et al., 2019). Notably different from other observations, is the extremely low abundance of the phylum Bacteroidetes in the current study. This may be due to a seasonal effect, as seasonality is a known phenomenon for this phylum (Díez-Vives et al., 2014) and there are studies showing that Bacteroidetes decrease significantly towards summer (Suh et al., 2015).

Compared to the algal surfaces, BF and SW generally showed slightly higher bacterial diversity than the algal surfaces, which further corroborates findings of multiple studies postulating that algae actively selects their associated microbiota. The only bacterial phylum represented on all algal surfaces, significantly abundant on FS, but not in the SW or BF reference samples was Deinococcota. They are known to be differentially abundant in different geographic regions and on seaweeds as, they play important role in denitrification and biofilm formation (Zulkifly et al., 2012; Wang et al., 2017; Pei et al., 2021; Qu et al., 2021). A member of the phylum is a known producer of the carotenoid deinoxanthin with known algicidal activity against dinoflagellates (Li et al., 2015), thus, members of this phylum may have important beneficial effects for the seaweed. The alphaproteobacterial genus *Octadecabacter* and members of the actinobacterial Sva0996 marine group were identified as differentially abundant on FV by LefSe analysis, but their ecological roles remain elusive as well as the role of the differentially abundant genus *Robiginitomaculum* on FE. The genus *Yoonia-Loktanella*, also significantly abundant on FE, is known for its ability to utilize complex algal exudates (Stock et al., 2022). It is a dimethylsulfopropionate (DMSP) degrader (Sun et al., 2020) with algicidal potential (Zheng et al., 2019), thus may be an important algal associate. Although all three algae were collected from the same spot, their microhabitats differ. FV in the Kiel Fjord grows close to the surface, with occasional desiccation events during unfavorable wind conditions whereas FS and FE occur at greater depths. In an earlier report on epibiosis at different seawater depths of 1-6m, the epibiont mass on FV was lowest at 6m (Rohde et al., 2008). Rohde et al. (2008) asserted that, the alga had to actively control fouling on its surface to overcome the stress of diminishing light intensity which comes with increasing depth in order to avoid the shading effect of the epibionts. This may be true for our current study as FV (shallow) was more fouled than FE and FS (deep).

Most published studies focus on the bacterial epiphytic community. A recent study however suggests to take eukaryotic diversity into account for explanation of prokaryotic community composition and dynamics in aquatic habitats (Bengtsson et al., 2017). In this study, ciliates were detected as most abundant eukaryotes based on ITS analyses except for the BF reference where Ascomycota were most abundant. Ciliates play key roles in the microbial food web (Chen et al., 2020). They are known bacterial grazers, hence generally influence bacterial epibiosis. However, the ciliate genus *Zoothamnium*, which was detected in high abundances on FV is also known as a marine pathogen (Shinn et al., 2015) and may contribute in part to the highly fouled surface observed. Significantly abundant ascomycete genus on FV was the yeast *Candida*, which might contribute to microfouling and consume algal exudates. The Ichthyosporea represents a fungus-like protozoan lineage not further classified. Their members including the genus *Sphaeroforma* which showed high abundances on FS surface have been previously detected in marine invertebrates (Jøstensen et al., 2002) but their role is as of yet unclear. *Alternaria*, which was second-most abundant fungal genus on FS and FE but only constituting 1.6% of the fungal community on FV, is known as an algal epiphyte known for producing compounds e.g. terpenes active against bacterial pathogens of algae (Shi et al., 2017). Sporidiobolales was significantly abundant on FE by LEfSe analysis. A member of this taxon is considered an antagonistic yeast with profound biocontrol efficiency against pathogens and may have positive relevance on the surface of FE (Dhanasekaran et al., 2021). Although the genus *Mucor* is generally known as endophytic or soil-derived, it has been also isolated from red algae (Dewapriya et al., 2014). Its main secondary metabolite was identified as tyrosol that displays antibiofilm activities in medical settings (Arias et al., 2016) and may exert similar ecological role on FE surface.

Unfortunately, not many microbial metabolites were putatively identified in this study although the surface extracts represent the combined surface metabolome of *Fucus* spp. and its associated epibionts. In the previous work, we annotated many microbial secondary metabolites in the surface extract of FV (Parrot et al., 2019). This may be due to factors such as seawater temperature, weather and seasons, which tend to influence surface metabolomes of seaweeds (Paix et al., 2019). In the current study, unannotated nodes in the MN may represent unidentified microbial compounds while microbial compounds in minute amounts may have be excluded during data pre-processing.

An earlier study reported FE to be less fouled than FV, after comparing the biomass and variation of their epiphytic community composition (Wikström and Kautsky, 2004). The low epiphytic growth on FE was partly attributed to its morphology, i.e., smoother surface of its fronds, thickness of cell-wall and surface texture (Wikström and Kautsky, 2004). Rickert et al. (2015) also reported a lesser seasonal epiphyte recruitment pattern in FE than FS. It is therefore conceivable, that in addition to the surface metabolite and microbiome, the thallus morphology of FE makes it a less suitable substratum for epibiosis, hence the observed least fouling state relative to the other *Fucus* spp.

## 5. Conclusion

In conclusion, we comparatively analyzed the surface metabolome and the epibiome of three co-occurring *Fucus* spp.; *F. distichus* subsp. *evanescens* (FE), *F. vesiculosus* (FV) and *F. serratus;* to gain insights into the potential factors underlying the observed differential fouling levels. The surface metabolites were clearly segregated from the tissue metabolites with many common metabolites among the three spp. FBMN and multivariate statistical analyses revealed many species-specific and discriminant surface metabolites such as MGTAs, DGTAs and carboxylic acid derivatives contributing to chemical differences on the algal surfaces. This study also provides evidence that the *Fucus* spp. differentially harbor diverse epiphytic prokaryotic and eukaryotic community influencing the surface biofilm. The species-specific and discriminant metabolites such as ulvaline, MGTA 20:5, MGTA 20:4, 4-pyridoxic acid, taxa (*Yoonia-Loktanella, Alternaria* and Mucoromycota) and the thallus morphology of FE, may directly and/or indirectly contribute to the least fouled surface observation. Our results show the ecological importance of some epibionts and surface metabolites on *Fucus* spp. and also have implications for the development of antifouling strategies and the conservation of coastal habitats.

## Abbreviations

*FE*: *Fucus distichus* subsp. *Evanescens*
FV: Fucus vesiculosus
FS: Fucus serratus
*SA*: surface adsorption
*SD*: solvent dipping
*SFA*: surface-free after surface adsorption
*SFD*: surface-free after solvent dipping
*W*: whole alga
*SW*: seawater
*BF*: stone biofilm

## Data availability

Raw reads from sequencing of 16S rDNA V3-V4 region amplicons and ITS2 amplicons have been deposited in the Sequence Read Archive (SRA) of National Center for Biotechnology Information (NCBI) database (BioProject: PRJNA992394). The MS data used for the molecular networking analysis were deposited in the MassIVE Public GNPS database under the accession number MSV000092347. The molecular networking job can be publicly accessed at https://gnps.ucsd.edu/ProteoSAFe/status.jsp?task=b56a28db5d8f4199b495e585c454a529 (accessed on 20 June, 2023)

## Acknowledgment

Arlette Wenzel-Storjohann is duly acknowledged for performing bioassays and her assistance during sampling.

## CRediT author statement

**Ernest Oppong-Danquah**: Methodology, Investigation, Formal analysis, Data curation, Visualization, Writing - Original Draft, Writing - Review & Editing. **Martina Blümel**: Methodology, Formal analysis, Visualization, Writing - Original Draft, Writing - Review & Editing, Visualization. **Deniz Tasdemir**: Conceptualization, Methodology, supervision, Writing - Review & Editing.

## References

Abe, S., Kaneda, T., 1975. Studies on the effect of marine products on cholesterol metabolism in rat-XI, isolation of a new betaine, ulvaline, from a green laver *Monostroma nitidum* and Its depressing effect on plasma cholesterol levels. Bull. Jap. Soc. Sci. Fish. 41, 567–571. https://doi.org/10.2331/suisan.41.567.

Alagawany, M., Elnesr, S.S., Farag, M.R., El-Naggar, K., Taha, A.E., Khafaga, A.F., et al., 2022. Betaine and related compounds: chemistry, metabolism and role in mitigating heat stress in poultry. J. Therm. Biol. 104, 103168. https://doi.org/10.1016/j.jtherbio.2021.103168.

Arias, L.S., Delbem, A.C., Fernandes, R.A., Barbosa, D.B., Monteiro, D.R., 2016. Activity of tyrosol against single and mixed-species oral biofilms. J. Appl. Microbiol. 120, 1240–9. https://doi.org/10.1111/jam.13070.

Bengtsson, M.M., Bühler, A., Brauer, A., Dahlke, S., Schubert, H., Blindow, I., 2017. Eelgrass leaf surface microbiomes are locally variable and highly correlated with epibiotic eukaryotes. Front. Microbiol. 8, 1–11. https://doi.org/10.3389/fmicb.2017.01312.

Bondoso, J., Balagué, V., Gasol, J.M., Lage, O.M., 2014. Community composition of the Planctomycetes associated with different macroalgae. FEMS Microbiol. Ecol. 88, 445–456. https://doi.org/10.1111/1574-6941.12258.

Bonilla-Rosso, G., Peimbert, M., Alcaraz, L.D., Hernández, I., Eguiarte, L.E., Olmedo-Alvarez, G., et al., 2012. Comparative metagenomics of two microbial mats at Cuatro Ciénegas Basin II: community structure and composition in oligotrophic environments. Astrobiology 12, 659–673. https://doi.org/10.1089/ast.2011.0724.

Buedenbender, L., Astone, F.A., Tasdemir, D., 2020. Bioactive molecular networking for mapping the antimicrobial constituents of the Baltic brown alga *Fucus vesiculosus*. Mar. Drugs 18, 1–21. https://doi.org/10.3390/md18060311.

Busetti, A., Maggs, C.A., Gilmore, B.F., 2017. Marine macroalgae and their associated microbiomes as a source of antimicrobial chemical diversity. Eur. J. Phycol. 52, 452–465. https://doi.org/10.1080/09670262.2017.1376709.

Callahan, B.J., McMurdie, P.J., Rosen, M.J., Han, A.W., Johnson, A.J.A., Holmes, S.P., 2016. DADA2: high resolution sample inference from Illumina amplicon data. Nat. Methods 13, 581–583. https://doi.org/10.1038/nmeth.3869.

Capistrant-Fossa, K.A., Morrison, H.G., Engelen, A.H., Quigley, C.T.C., Morozov, A., Serrão, E.A., et al., 2021. The microbiome of the habitat-forming brown alga *Fucus vesiculosus* (Phaeophyceae) has similar cross-Atlantic structure that reflects past and present drivers1. J. Phycol. 57, 1681–1698. https://doi.org/10.1111/jpy.13194.

Chambers, M.C., Maclean, B., Burke, R., Amodei, D., Ruderman, D.L., Neumann, S., et al., 2012. A cross platform toolkit for mass spectrometry and proteomics. Nat. Biotechnol. 30, 918–920. https://doi.org/10.1038/nbt.2377.

Chen, W.-L., Chiang, K.-P., Tsai, S.-F., 2020. Neglect of presence of bacteria leads to inaccurate growth parameters of the oligotrich ciliate *Strombidium* sp. during grazing experiments on nanoflagellates. Front. Mar. Sci. 7, 1–10. https://doi.org/10.3389/fmars.2020.569309.

Cirri, E., Grosser, K., Pohnert, G., 2016. A solid phase extraction based non-disruptive sampling technique to investigate the surface chemistry of macroalgae. Biofouling 32, 145–153. https://doi.org/10.1080/08927014.2015.1130823.

Corrigan, S., Brown, A.R., Tyler, C.R., Wilding, C., Daniels, C., Ashton, I.G.C., et al., 2023. Development and diversity of epibiont assemblages on cultivated sugar Kelp (*Saccharina latissima*) in relation to farming schedules and harvesting techniques. Life 13, 1–22. https://doi.org/10.3390/life13010209.

Coyer, J., Peters, A., Hoarau, G., Stam, W., Olsen, J., 2002. Hybridization of the marine seaweeds, *Fucus serratus* and *Fucus evanescens* (Heterokontophyta: Phaeophyceae) in a 100-year-old zone of secondary contact. Proc. R. Soc. Lond., Ser. B: Biol. Sci. 269, 1829–1834. https://doi.org/10.1098/rspb.2002.2093.

Creed, J.C., Norton, T.A., Kain, J.M., 1997. Intraspecific competition in *Fucus serratus* germlings: the interaction of light, nutrients and density. J. Exp. Mar. Biol. Ecol. 212, 211–223. https://doi.org/10.1016/S0022-0981(96)02748-7.

Croft, M.T., Lawrence, A.D., Raux-Deery, E., Warren, M.J., Smith, A.G., 2005. Algae acquire vitamin B12 through a symbiotic relationship with bacteria. Nature 438, 90–93. https://doi.org/10.1038/nature04056.

da Gama, B.A.P., Plouguerné, E., Pereira, R.C., 2014. The antifouling defence mechanisms of marine macroalgae. Adv. Bot. Res. 71, 413–440. https://doi.org/10.1016/B978-0-12-408062-1.00014-7.

da Silva, J.C., Lombardi, A.T., 2020. Chlorophylls in microalgae: occurrence, distribution, and biosynthesis. In: Jacob-Lopes, E., Queiroz, M.I., Zepka, L.Q., (Eds.), Pigments from Microalgae Handbook. Springer International Publishing, Cham, pp. 1–18.

da Silva, R.R., Wang, M., Nothias, L.-F., van der Hooft, J.J.J., Caraballo-Rodríguez, A.M., Fox, E., et al., 2018. Propagating annotations of molecular networks using *in silico* fragmentation. PLoS Comp. Biol. 14, 1–26. https://doi.org/10.1371/journal.pcbi.1006089.

Dewapriya, P., Li, Y.-X., Himaya, S.W.A., Kim, S.-K., 2014. Isolation and characterization of marine derived *Mucor* sp. for the fermentative production of tyrosol. Process Biochem. 49, 1402–1408. https://doi.org/10.1016/j.procbio.2014.06.004.

Dhanasekaran, S., Yang, Q., Godana, E.A., Liu, J., Li, J., Zhang, H., 2021. Trehalose supplementation enhanced the biocontrol efficiency of *Sporidiobolus pararoseus* Y16 through increased oxidative stress tolerance and altered transcriptome. Pest Manag. Sci. 77, 4425–4436. https://doi.org/10.1002/ps.6477.

Díez-Vives, C., Gasol, J.M., Acinas, S.G., 2014. Spatial and temporal variability among marine Bacteroidetes populations in the NW Mediterranean sea. Syst. Appl. Microbiol. 37, 68–78. https://doi.org/10.1016/j.syapm.2013.08.006.

Dittami, S.M., Gravot, A., Renault, D., Goulitquer, S., Eggert, A., Bouchereau, A., et al., 2011. Integrative analysis of metabolite and transcript abundance during the short-term response to saline and oxidative stress in the brown alga *Ectocarpus siliculosus*. Plant, Cell Environ. 34, 629–642. https://doi.org/10.1111/j.1365-3040.2010.02268.x.

Egan, S., Harder, T., Burke, C., Steinberg, P., Kjelleberg, S., Thomas, T., 2013. The seaweed holobiont: understanding seaweed–bacteria interactions. FEMS Microbiol. Rev. 37, 462–476. https://doi.org/10.1111/1574-6976.12011.

Egan, S., Thomas, T., Kjelleberg, S., 2008. Unlocking the diversity and biotechnological potential of marine surface associated microbial communities. Curr. Opin. Microbiol. 11, 219–225. https://doi.org/10.1016/j.mib.2008.04.001.

El-Manaway, I.M., Rashedy, S.H., 2022. The ecology and physiology of seaweeds: an overview. In: Ranga Rao, A., Ravishankar, G.A., (Eds.), Sustainable Global Resources Of Seaweeds Volume 1: Bioresources, cultivation, trade and multifarious applications. 1. Springer International Publishing, Cham, pp. 3–16.

Ernst, M., Kang, K.B., Caraballo-Rodríguez, A.M., Nothias, L.-F., Wandy, J., Chen, C., et al., 2019. MolNetEnhancer: enhanced molecular networks by integrating metabolome mining and annotation tools. Metabolites 9, 1–25. https://doi.org/10.3390/metabo9070144.

Feunang, Y.D., Eisner, R., Knox, C., Chepelev, L., Hastings, J., Owen, G., et al., 2016. ClassyFire: automated chemical classification with a comprehensive, computable taxonomy. Journal of Cheminformatics 8, 1–20. https://doi.org/10.1186/s13321-016-0174-y.

Filion-Myklebust, C., 1981. Epidermis shedding in the brown seaweed *Ascophyllum nodosum* (L.) Le Jolis and its ecological significance. Mar. Biol. Lett. 2, 45–51.

Givskov, M., de Nys, R., Manefield, M., Gram, L., Maximilien, R., Eberl, L., et al., 1996. Eukaryotic interference with homoserine lactone-mediated prokaryotic signalling. J. Bacteriol. 178, 6618–6622. https://doi.org/10.1128/jb.178.22.6618-6622.1996.

Grant, M.A.A., Kazamia, E., Cicuta, P., Smith, A.G., 2014. Direct exchange of vitamin B12 is demonstrated by modelling the growth dynamics of algal–bacterial cocultures. ISME J. 8, 1418–1427. https://doi.org/10.1038/ismej.2014.9.

Grinberg, M., Orevi, T., Kashtan, N., 2019. Bacterial surface colonization, preferential attachment and fitness under periodic stress. PLoS Comp. Biol. 15, e1006815. https://doi.org/10.1371/journal.pcbi.1006815.

Groisillier, A., Shao, Z., Michel, G., Goulitquer, S., Bonin, P., Krahulec, S., et al., 2013. Mannitol metabolism in brown algae involves a new phosphatase family. J. Exp. Bot. 65, 559–570. https://doi.org/10.1093/jxb/ert405.

Harder, T., 2009. Marine epibiosis: concepts, ecological consequences and host defence. In: Flemming, H., Murthy, P.S., Venkatesan, R., Cooksey, K., (Eds.), Marine and Industrial Biofouling. Springer, pp. 219–231.

Heavisides, E., Rouger, C., Reichel, A., Ulrich, C., Wenzel-Storjohann, A., Sebens, S., et al., 2018. Seasonal variations in the metabolome and bioactivity profile of *Fucus vesiculosus* extracted by an optimised, pressurised liquid extraction protocol. Mar. Drugs 16, 1–28. https://doi.org/10.3390/md16120503.

Hoffmann, D.Y., Shachar-Hill, Y., 2023. Do betaine lipids replace phosphatidylcholine as fatty acid editing hubs in microalgae? Front. Plant Sci. 14, 1–14. https://doi.org/10.3389/fpls.2023.1077347.

Honkanen, T., Jormalainen, V., 2005. Genotypic variation in tolerance and resistance to fouling in the brown alga *Fucus vesiculosus*. Oecologia 144, 196–205. https://doi.org/10.1007/s00442-005-0053-0.

Horai, H., Arita, M., Kanaya, S., Nihei, Y., Ikeda, T., Suwa, K., et al., 2010. MassBank: a public repository for sharing mass spectral data for life sciences. J. Mass Spectrom. 45, 703–714. https://doi.org/10.1002/jms.1777.

Jøstensen, J.-P., Sperstad, S., Johansen, S., Landfald, B., 2002. Molecular-phylogenetic, structural and biochemical features of a cold-adapted, marine ichthyosporean near the animal-fungal divergence, described from in vitro cultures. Eur. J. Protistol. 38, 93–104. https://doi.org/10.1078/0932-4739-00855.

Koivikko, R., Loponen, J., Honkanen, T., Jormalainen, V., 2005. Contents of soluble, cell-wall-bound and exuded phlorotanins in the brown alga *Fucus vesiculosus*, with implications on the their ecological functions. J. Chem. Ecol. 31, 195–212. https://doi.org/10.1007/s10886-005-0984-2.

Kolanjinathan, K., Ganesh, P., Saranraj, P., 2014. Pharmacological importance of seaweeds: a review. World J. Fish. Mar. Sci. 6, 1–15. https://doi.org/10.5829/idosi.wjfms.2014.06.01.76195.

Krause-Jensen, D., Duarte, C.M., 2016. Substantial role of macroalgae in marine carbon sequestration. Nature Geoscience 9, 737–742. https://doi.org/10.1038/ngeo2790.

Lachnit, T., Fischer, M., Künzel, S., Baines, J.F., Harder, T., 2013. Compounds associated with algal surfaces mediate epiphytic colonization of the marine macroalga *Fucus vesiculosus*. FEMS Microbiol. Ecol. 84, 411–420. https://doi.org/10.1111/1574-6941.12071.

Lachnit, T., Meske, D., Wahl, M., Harder, T., Schmitz, R., 2011. Epibacterial community patterns on marine macroalgae are host-specific but temporally variable. Environ. Microbiol. 13, 655–665. https://doi.org/10.1111/j.1462-2920.2010.02371.x.

Lachnit, T., Wahl, M., Harder, T., 2010. Isolated thallus-associated compounds from the macroalga *Fucus vesiculosus* mediate bacterial surface colonization in the field similar to that on the natural alga. Biofouling 26, 247–255. https://doi.org/10.1080/08927010903474189.

Latorre, N., Castañeda, F., Meynard, A., Rivas, J., Contreras-Porcia, L., 2019. First approach of characterization of bioactive compound in *Pyropia orbicularis* during the daily tidal cycle. Lat. Am. J. Aquat. Res. 47, 826–840. https://doi.org/10.3856/vol47-issue5-fulltext-12.

Li, Y., Zhu, H., Lei, X., Zhang, H., Guan, C., Chen, Z., et al., 2015. The first evidence of deinoxanthin from *Deinococcus* sp. Y35 with strong algicidal effect on the toxic dinoflagellate *Alexandrium tamarense*. J. Hazard. Mater. 290, 87–95. https://doi.org/10.1016/j.jhazmat.2015.02.070.

Littler, M.M., Littler, D.S., 1999. Epithallus sloughing: a self-cleaning mechanism for coralline algae. Coral Reefs 18, 204. https://doi.org/10.1007/s003380050182.

Liu, C., Cui, Y., Li, X., Yao, M., 2021. *microeco*: an R package for data mining in microbial community ecology. FEMS Microbiol. Ecol. 97, fiaa255. https://doi.org/10.1093/femsec/fiaa255.

Llewellyn, C.A., Sommer, U., Dupont, C.L., Allen, A.E., Viant, M.R., 2015. Using community metabolomics as a new approach to discriminate marine microbial particulate organic matter in the western English Channel. Prog. Oceanogr. 137, 421–433. https://doi.org/10.1016/j.pocean.2015.04.022.

Magoč, T., Salzberg, S.L., 2011. FLASH: fast length adjustment of short reads to improve genome assemblies. Bioinformatics 27, 2957–2963. https://doi.org/10.1093/bioinformatics/btr507.

McArthur, D.M., Moss, B.L., 1977. The ultrastructure of cell walls in *Enteromorpha intestinalis* (L.) link. Br. Phycol. J. 12, 359–368. https://doi.org/10.1080/00071617700650381.

McMurdie, P.J., Holmes, S., 2013. phyloseq: an R package for reproducible interactive analysis and graphics of microbiome census data. PloS one 8, e61217. https://doi.org/10.1371/journal.pone.0061217.

Mensch, B., Neulinger, S.C., Graiff, A., Pansch, A., Künzel, S., Fischer, M.A., et al., 2016. Restructuring of epibacterial communities on *Fucus vesiculosus* forma *mytili* in response to elevated *p*CO2 and increased temperature levels. Front. Microbiol. 7, 1–15. https://doi.org/10.3389/fmicb.2016.00434.

Menshova, R.V., Shevchenko, N.M., Imbs, T.I., Zvyagintseva, T.N., Malyarenko, O.S., Zaporoshets, T.S., et al., 2016. Fucoidans from brown alga *Fucus evanescens*: structure and biological activity. Front. Mar. Sci. 3, 129. https://doi.org/10.3389/fmars.2016.00129.

Mohimani, H., Gurevich, A., Shlemov, A., Mikheenko, A., Korobeynikov, A., Cao, L., et al., 2018. Dereplication of microbial metabolites through database search of mass spectra. Nat. Commun. 9, 4035. https://doi.org/10.1038/s41467-018-06082-8.

Monds, R.D., O’Toole, G.A., 2009. The developmental model of microbial biofilms: ten years of a paradigm up for review. Trends Microbiol. 17, 73–87. https://doi.org/10.1016/j.tim.2008.11.001.

Murali, A., Bhargava, A., Wright, E.S., 2018. IDTAXA: a novel approach for accurate taxonomic classification of microbiome sequences. Microbiome 6, 1–14. https://doi.org/10.1186/s40168-018-0521-5.

Negara, B.F.S.P., Sohn, J.-H., Kim, J.-S., Choi, J.-S., 2021. Antifungal and larvicidal activities of phlorotannins from brown seaweeds. Mar. Drugs 19, 223. https://doi.org/10.3390/md19040223.

Noisette, F., Hurd, C., 2018. Abiotic and biotic interactions in the diffusive boundary layer of kelp blades create a potential refuge from ocean acidification. Funct. Ecol. 32, 1329–1342. https://doi.org/10.1111/1365-2435.13067.

Noorjahan, A., Mahesh, S., Aiyamperumal, B., Anantharaman, P., 2022. Exploring Marine Fungal Diversity and Their Applications in Agriculture. In: Rajpal, V.R., Singh, I., Navi, S.S., (Eds.), Fungal diversity, ecology and control management. Springer Nature Singapore, Singapore, pp. 293–310.

Nothias, L.-F., Petras, D., Schmid, R., Dührkop, K., Rainer, J., Sarvepalli, A., et al., 2020. Feature-based molecular networking in the GNPS analysis environment. Nat. Methods 17, 905–908. https://doi.org/10.1038/s41592-020-0933-6.

Nylund, G.M., Cervin, G., Persson, F., Hermansson, M., Steinberg, P.D., Pavia, H., 2008. Seaweed defence against bacteria: a poly-brominated 2-heptanone from the red alga *Bonnemaisonia hamifera* inhibits bacterial colonisation. Mar. Ecol. Prog. Ser. 369, 39–50. https://doi.org/10.3354/meps07577.

Okeke, E.S., Nweze, E.J., Chibuogwu, C.C., Anaduaka, E.G., Chukwudozie, K.I., Ezeorba, T.P.C., 2021. Aquatic phlorotannins and human health: bioavailability, toxicity, and future prospects. Nat. Prod. Commun. 16, 1–23. https://doi.org/10.1177/1934578x211056144.

Ould, E., Caldwell, G.S., 2022. The potential of seaweed for carbon capture. CABI Rev. 17, 009. https://doi.org/10.1079/cabireviews202217009.

Paix, B., Othmani, A., Debroas, D., Culioli, G., Briand, J.-F., 2019. Temporal covariation of epibacterial community and surface metabolome in the Mediterranean seaweed holobiont *Taonia atomaria*. Environ. Microbiol. 21, 3346–3363. https://doi.org/10.1111/1462-2920.14617.

Pang, Z., Chong, J., Zhou, G., de Lima Morais, D.A., Chang, L., Barrette, M., et al., 2021. MetaboAnalyst 5.0: narrowing the gap between raw spectra and functional insights. Nucleic Acids Res. 49, W388–W396. https://doi.org/10.1093/nar/gkab382.

Pantos, O., Bongaerts, P., Dennis, P.G., Tyson, G.W., Hoegh-Guldberg, O., 2015. Habitat-specific environmental conditions primarily control the microbiomes of the coral *Seriatopora hystrix*. ISME J. 9, 1916–1927. https://doi.org/10.1038/ismej.2015.3.

Papazian, S., Parrot, D., Burýšková, B., Weinberger, F., Tasdemir, D., 2019. Surface chemical defence of the eelgrass *Zostera marina* against microbial foulers. Sci. Rep. 9, 3323. https://doi.org/10.1038/s41598-019-39212-3.

Parrot, D., Blümel, M., Utermann, C., Chianese, G., Krause, S., Kovalev, A., et al., 2019. Mapping the surface microbiome and metabolome of brown seaweed *Fucus vesiculosus* by amplicon sequencing, integrated metabolomics and imaging techniques. Sci. Rep. 9, 1061. https://doi.org/10.1038/s41598-018-37914-8.

Pei, P., Aslam, M., Du, H., Liang, H., Wang, H., Liu, X., et al., 2021. Environmental factors shape the epiphytic bacterial communities of *Gracilariopsis lemaneiformis*. Sci. Rep. 11, 8671. https://doi.org/10.1038/s41598-021-87977-3.

Pendyala, M., Parvataneni, R., Krishna, N., Rao, D., Rao, C., 2003. Sphingolipids from marine organisms: a review. Nat. Prod. Sci. 9, 1–27.

Plouguerné, E., Ioannou, E., Georgantea, P., Vagias, C., Roussis, V., Hellio, C., et al., 2010. Anti microfouling activity of lipidic metabolites from the invasive brown alga *Sargassum muticum* (Yendo) Fensholt. Mar. Biotechnol. 12, 52–61. https://doi.org/10.1007/s10126-009-9199-9.

Plouguerné, E., Souza, L., Sassaki, G., Hellio, C., Trepos, R., Da Gama, B., et al., 2020. Glycoglycerolipids from *Sargassum vulgare* as potential antifouling agents. Front. Mar. Sci. 7. https://doi.org/10.3389/fmars.2020.00116.

Pluskal, T., Castillo, S., Villar-Briones, A., Oresic, M., 2010. MZmine 2: modular framework for processing, visualizing, and analyzing mass spectrometry-based molecular profile data. BMC Bioinformatics 11, 1–11. https://doi.org/10.1186/1471-2105-11-395.

Potin, P., 2008. Oxidative burst and related responses in biotic interactions of algae. In: Amsler, C.D., (Ed.), Algal Chemical Ecology. Springer, pp. 245–271.

Qu, J., Yang, H., Liu, Y., Qi, H., Wang, Y., Zhang, Q., 2021. The study of natural biofilm formation and microbial community structure for recirculating aquaculture system. IOP Conf. Ser.: Earth Environ. Sci. 742, 012018. https://doi.org/10.1088/1755-1315/742/1/012018.

Quémener, M., Kikionis, S., Fauchon, M., Toueix, Y., Aulanier, F., Makris, A.M., et al., 2022. Antifouling activity of halogenated compounds derived from the red alga *Sphaerococcus coronopifolius*: potential for the development of environmentally friendly solutions. Mar. Drugs 20, 1–16. https://doi.org/10.3390/md20010032.

Quigley, C.T.C., Capistrant-Fossa, K.A., Morrison, H.G., Johnson, L.E., Morozov, A., Hertzberg, V.S., et al., 2020. Bacterial communities show algal host (*Fucus* spp.)/zone differentiation across the stress gradient of the intertidal zone. Front. Microbiol. 11, 563118. https://doi.org/10.3389/fmicb.2020.563118.

Rickert, E., Karsten, U., Pohnert, G., Wahl, M., 2015. Seasonal fluctuations in chemical defenses against macrofouling in *Fucus vesiculosus* and *Fucus serratus* from the Baltic Sea. Biofouling 31, 363–377. https://doi.org/10.1080/08927014.2015.1041020.

Rickert, E., Lenz, M., Barboza, F.R., Gorb, S.N., Wahl, M., 2016a. Seasonally fluctuating chemical microfouling control in *Fucus vesiculosus* and *Fucus serratus* from the Baltic Sea. Mar. Biol. 163, 203. https://doi.org/10.1007/s00227-016-2970-3.

Rickert, E., Wahl, M., Link, H., Richter, H., Pohnert, G., 2016b. Seasonal variations in surface metabolite composition of *Fucus vesiculosus* and *Fucus serratus* from the Baltic Sea. PLOS ONE 11, e0168196. https://doi.org/10.1371/journal.pone.0168196.

Rohde, S., Hiebenthal, C., Wahl, M., Karez, R., Bischof, K., 2008. Decreased depth distribution of *Fucus vesiculosus* (Phaeophyceae) in the Western Baltic: effects of light deficiency and epibionts on growth and photosynthesis. Eur. J. Phycol. 43, 143–150. https://doi.org/10.1080/09670260801901018.

Sachindra, N.M., Sato, E., Maeda, H., Hosokawa, M., Niwano, Y., Kohno, M., et al., 2007. Radical scavenging and singlet oxygen quenching activity of marine carotenoid fucoxanthin and its metabolites. J. Agric. Food Chem. 55, 8516–8522. https://doi.org/10.1021/jf071848a.

Saha, M., Goecke, F., Bhadury, P., 2018. Minireview: algal natural compounds and extracts as antifoulants. J. Appl. Phycol. 30, 1875–1875. https://doi.org/10.1007/s10811-018-1424-3.

Saha, M., Rempt, M., Grosser, K., Pohnert, G., Weinberger, F., 2011. Surface-associated fucoxanthin mediates settlement of bacterial epiphytes on the rockweed *Fucus vesiculosus*. Biofouling 27, 423–433. https://doi.org/10.1080/08927014.2011.580841.

Selvarajan, R., Sibanda, T., Tekere, M., 2018. Thermophilic bacterial communities inhabiting the microbial mats of “indifferent” and chalybeate (iron-rich) thermal springs: diversity and biotechnological analysis. MicrobiologyOpen 7, e00560. https://doi.org/10.1002/mbo3.560.

Selvarajan, R., Sibanda, T., Venkatachalam, S., Ogola, H.J.O., Christopher Obieze, C., Msagati, T.A., 2019. Distribution, interaction and functional profiles of epiphytic bacterial communities from the rocky intertidal seaweeds, South Africa. Sci. Rep. 9, 1–13. https://doi.org/10.1038/s41598-019-56269-2.

Shannon, P., Markiel, A., Ozier, O., Baliga, N.S., Wang, J.T., Ramage, D., et al., 2003. Cytoscape: a software environment for integrated models of biomolecular interaction networks. Genome Res. 13, 2498–2504. https://doi.org/10.1101/gr.1239303.

Shi, Z.-Z., Miao, F.-P., Fang, S.-T., Liu, X.-H., Yin, X.-L., Ji, N.-Y., 2017. Sesteralterin and tricycloalterfurenes A–D: terpenes with rarely occurring frameworks from the marine-alga-epiphytic fungus *Alternaria alternata* k21-1. J. Nat. Prod. 80, 2524–2529. https://doi.org/10.1021/acs.jnatprod.7b00478.

Shinn, A.P., Mühlhölzl, A.P., Coates, C.J., Metochis, C., Freeman, M.A., 2015. Zoothamnium duplicatum infestation of cultured horseshoe crabs (*Limulus polyphemus*). J. Invertebr. Pathol. 125, 81–86. https://doi.org/10.1016/j.jip.2014.12.002.

Singh, B., 2014. Antibacterial activity of glycerol, lactose, maltose, mannitol, raffinose and xylose. Noto are Medicine 17223318.

Sofy, A.R., Dawoud, R.A., Sofy, M.R., Mohamed, H.I., Hmed, A.A., El-Dougdoug, N.K., 2020. Improving regulation of enzymatic and non-enzymatic antioxidants and stress-related gene stimulation in cucumber mosaic cucumovirus-infected cucumber plants treated with glycine betaine, chitosan and combination. Molecules 25, 2341. https://doi.org/10.3390/molecules25102341.

Soman, A.G., Gloer, J.B., Koster, B., Malloch, D., 1999. Sporovexins A− C and a new preussomerin analog: antibacterial and antifungal metabolites from the coprophilous fungus *Sporormiella vexans*. J. Nat. Prod. 62, 659–661. https://doi.org/10.1021/np980563c.

Stock, W., Willems, A., Mangelinckx, S., Vyverman, W., Sabbe, K., 2022. Selection constrains lottery assembly in the microbiomes of closely related diatom species. ISME Communications 2, 11. https://doi.org/10.1038/s43705-022-00091-x.

Stratil, S.B., Neulinger, S.C., Knecht, H., Friedrichs, A.K., Wahl, M., 2013. Temperature-driven shifts in the epibiotic bacterial community composition of the brown macroalga *Fucus vesiculosus*. MicrobiologyOpen 2, 338–349. https://doi.org/10.1002/mbo3.79.

Suh, S.-S., Park, M., Hwang, J., Kil, E.-J., Jung, S.W., Lee, S., et al., 2015. Seasonal dynamics of marine microbial community in the South Sea of Korea. PLOS ONE 10, e0131633. https://doi.org/10.1371/journal.pone.0131633.

Sun, H., Tan, S., Liang, J., Yang, P., Xin, Y., Zhang, X., 2020. Horizontal and vertical distribution of dimethylsulfoniopropionate (DMSP) producing and catabolizing bacteria in the East China Sea. Acta Pharm. Sin 60, 1865–1881.

Wahl, M., Goecke, F., Labes, A., Dobretsov, S., Weinberger, F., 2012. The second skin: ecological role of epibiotic biofilms on marine organisms. Front. Microbiol. 3, 292. https://doi.org/10.3389/fmicb.2012.00292.

Wahl, M., Shahnaz, L., Dobretsov, S., Saha, M., Symanowski, F., David, K., et al., 2010. Ecology of antifouling resistance in the bladder wrack *Fucus vesiculosus*: patterns of microfouling and antimicrobial protection. Mar. Ecol. Prog. Ser. 411, 33–48. https://doi.org/10.3354/meps08644.

Wang, M., Carver, J.J., Phelan, V.V., Sanchez, L.M., Garg, N., Peng, Y., et al., 2016. Sharing and community curation of mass spectrometry data with Global Natural Products Social Molecular Networking. Nat. Biotechnol. 34, 828–837. https://doi.org/10.1038/nbt.3597.

Wang, X.-X., Fang, F., Chen, Y.-P., Guo, J.-S., Li, K., Wang, H., 2017. N2O micro-profiles in biofilm from a one-stage autotrophic nitrogen removal system by microelectrode. Chemosphere 175, 482–489. https://doi.org/10.1016/j.chemosphere.2017.02.026.

Wheeler, G.L., Tait, K., Taylor, A., Brownlee, C., Joint, I., 2006. Acyl-homoserine lactones modulate the settlement rate of zoospores of the marine alga *Ulva intestinalis* via a novel chemokinetic mechanism. Plant, Cell Environ. 29, 608–618. https://doi.org/10.1111/j.1365-3040.2005.01440.x.

Wikström, S.A., Kautsky, L., 2004. Invasion of a habitat-forming seaweed: effects on associated biota. Biol. Invasions 6, 141–150. https://doi.org/10.1023/B:BINV.0000022132.00398.14.

Wikström, S.A., von Wachenfeldt, T., Kautsky, L., 2002. Establishment of the exotic species *Fucus evanescens* C. Ag. (Phaeophyceae) in Öresund, southern Sweden. Bot. Mar. 45, 510–517. https://doi.org/doi:10.1515/BOT.2002.054.

Wilson, M.P., Footitt, E.J., Papandreou, A., Uudelepp, M.-L., Pressler, R., Stevenson, D.C., et al., 2017. An LC–MS/MS-based method for the quantification of pyridox(am)ine 5′-phosphate oxidase activity in dried blood spots from patients with epilepsy. Anal. Chem. 89, 8892–8900. https://doi.org/10.1021/acs.analchem.7b01358.

Zheng, N., Sun, L., Ding, N., Li, C., Fu, B., Wang, C., et al., 2019. Diversity of algicidal bacteria associated with harmful microalgae and the algicidal mechanisms. Microbiol. China 46, 1204–1219.

Zulkifly, S., Hanshew, A., Young, E.B., Lee, P., Graham, M.E., Graham, M.E., et al., 2012. The epiphytic microbiota of the globally widespread macroalga *Cladophora glomerata* (Chlorophyta, Cladophorales). Am. J. Bot. 99, 1541–1552. https://doi.org/10.3732/ajb.1200161.

