## Supplemental Tables and Figures for "Metabolomics and Microbiomics Insights into the Differential Surface Fouling of Brown Algae"

List of Figures — Page No.

Figure S1: UPLC-MS(+) base peak chromatograms of Surface Adsorption (SA) extracts of A) FE, B) FS and C) FV — 4

Figure S2: UPLC-MS(+) base peak chromatograms of Solvent Dipping (SD) extracts of A) FE, B) FS and C) FV — 5

Figure S3: UPLC-MS(+) base peak chromatograms of Surface-free after adsorption (SFA) extracts of A) FE, B) FS and C) FV — 6

Figure S4: UPLC-MS(+) base peak chromatograms of Surface-free after dipping (SFD) extracts of A) FE, B) FS and C) FV — 7

Figure S5: UPLC-MS(+) base peak chromatograms of Whole (W) extracts of A) FE, B) FS and C) FV — 8

Figure S6: MN generated from UPLC-(-)-ESI-MS/MS data of all extracts of *Fucus* spp. in the negative ion mode — 9

Figure S7: Variation of the most significant discriminatory metabolite markers (VIP >1.8) on the surfaces of *Fucus* spp. — 10

Figure S8: PCA scores plot generated from UPLC-(+)-ESI-MS data of all extracts — 11

Figure S9: MN generated from UPLC-(+)-ESI-MS/MS data of all surface-free and whole extracts 12

Figure S10: Rarefaction curves of bacterial V3/V4 region amplicon sequences from all 66 samples. 13

Figure S11: Alpha diversity (ASV-Observed vs. Shannon) of bacterial epiphytic community with regard to sample source 14

Figure S12: Alpha diversity (ASV-Observed vs. Shannon) of bacterial epiphytic community with regard to individual. 15

Figure S13: Beta diversity analysis of bacterial amplicon data based on Bray-Curtis distance calculation on *Fucus* spp. 16

Figure S14: Bacterial orders associated to surfaces of *Fucus* spp. 17

Figure S15: Rarefaction curves of eukaryotic ITS region amplicon sequences from all 60 samples. 18

Figure S16: Alpha diversity (ASV-Observed vs. Shannon) of eukaryotic epiphytic community based on ITS fragment sequences with regard to sample source. 19

Figure S17: Alpha diversity (ASV-Observed vs. Shannon) of eukaryotic epiphytic community based on ITS fragment sequences with regard to individual 20

Figure S18: Alpha diversity (ASV-Observed vs. Shannon) of fungal epiphytic community based on ITS fragment sequences with regard to sample source. 21

Figure S19: Alpha diversity (ASV-Observed vs. Shannon) of fungal epiphytic community based on ITS fragment sequences with regard to individual. 22

Figure S20: Fungal orders associated to surfaces of *Fucus* spp., stone biofilm (BF) and seawater (SW) reference samples. 23

List of Tables Page No.

Table S2: Relative abundances of bacterial genera (> 1%) associated to surfaces *Fucus* spp.,

Seawater and stone biofilm 24

Table S3: Bacterial beta diversity statistics based on Bray-Curtis dissimilarity 25

Table S4: Relative abundances of eukaryote genera (> 1%) associated to surfaces of *Fucus* spp., seawater and stone biofilm . 26

Table S5: ITS beta diversity statistics based on Bray-Curtis dissimilarity 27

Table S6: Relative abundances of fungal genera (>1%) associated to surfaces of *Fucus* sp., seawater and stone biofilm 28

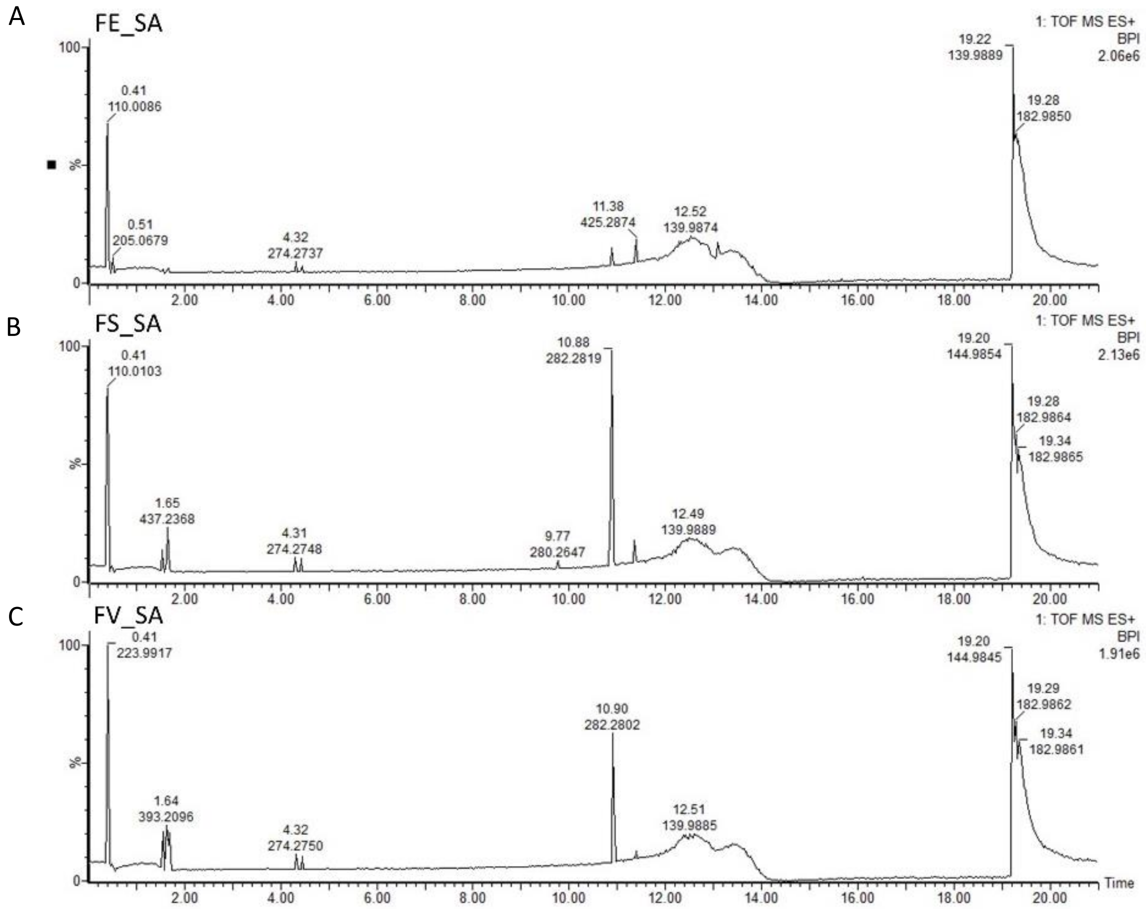


Figure S1: UPLC-MS(+) base peak chromatograms of Surface Adsorption (SA) extracts of A) FE, B) FS and C) FV

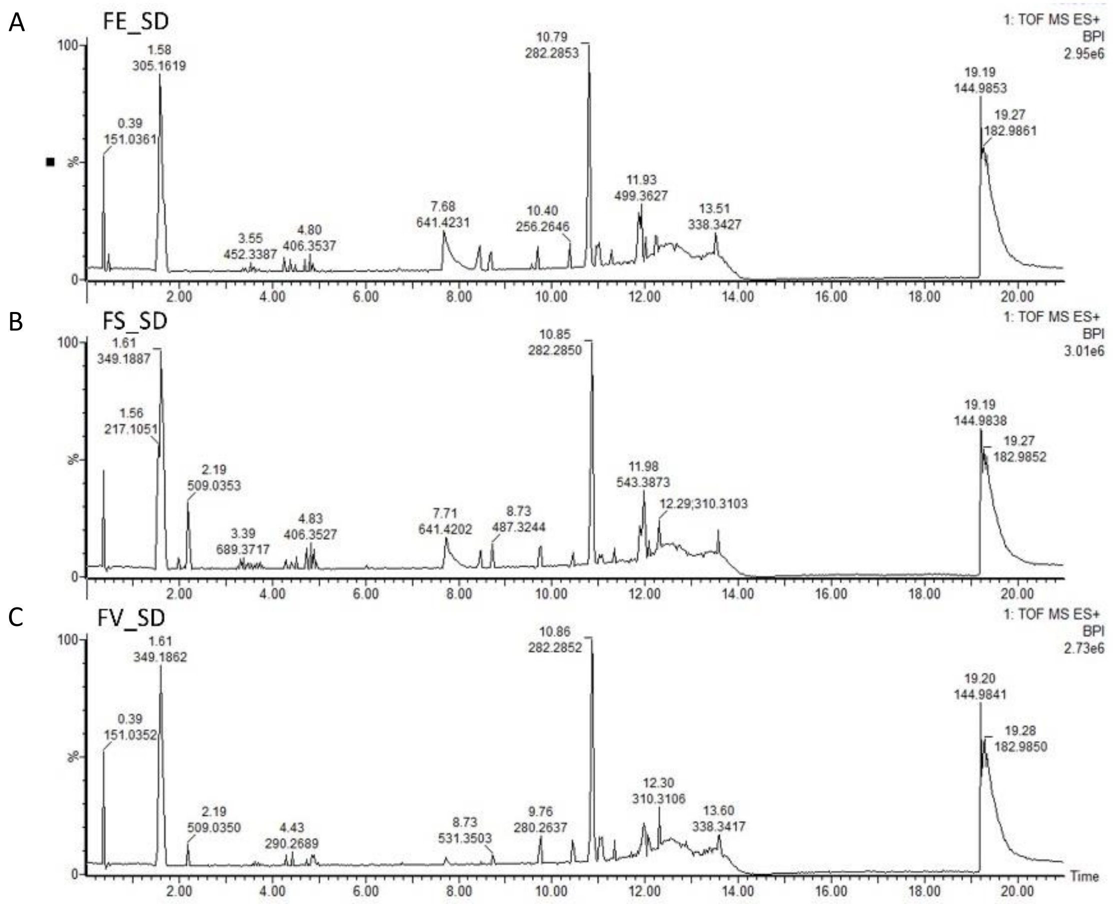


Figure S2: UPLC-MS(+) base peak chromatograms of Solvent Dipping (SD) extracts of A) FE, B) FS and C) FV

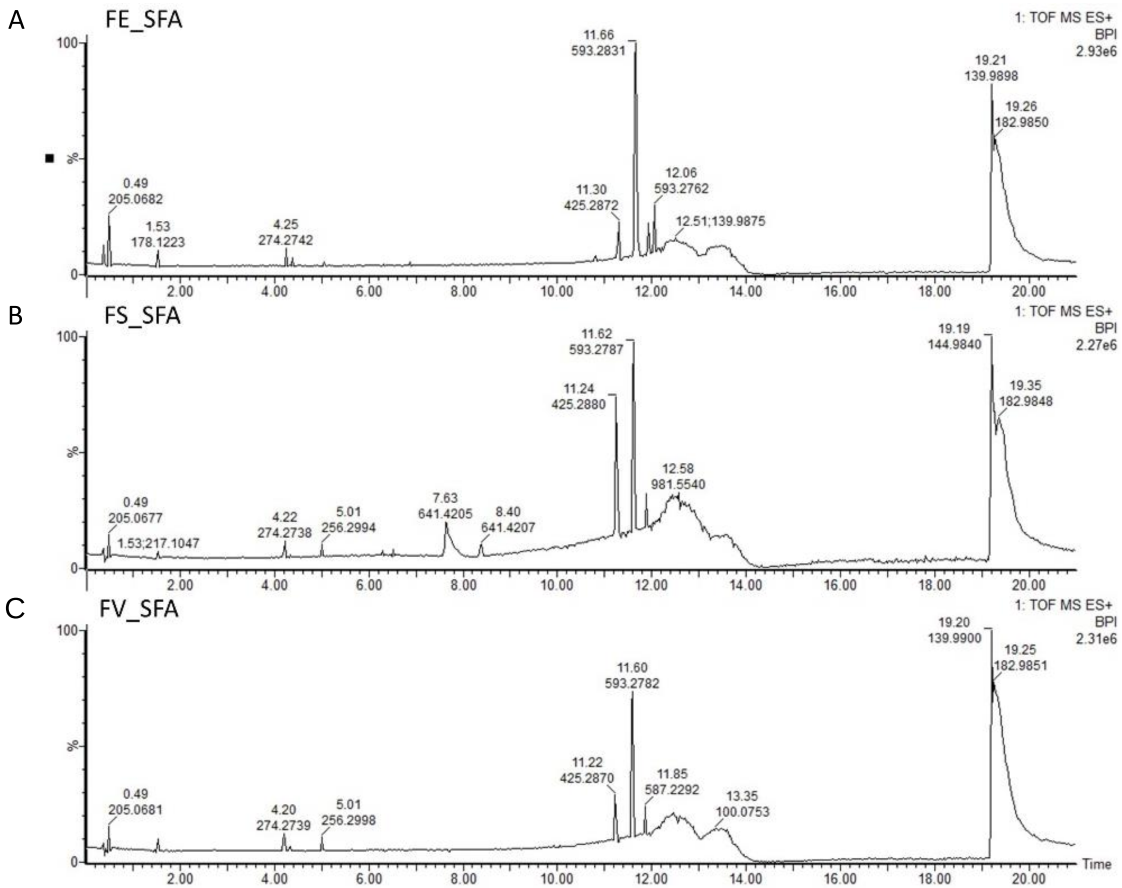


Figure S3: UPLC-MS(+) base peak chromatograms of Surface-free after adsorption (SFA) extracts of A) FE, B) FS and C) FV

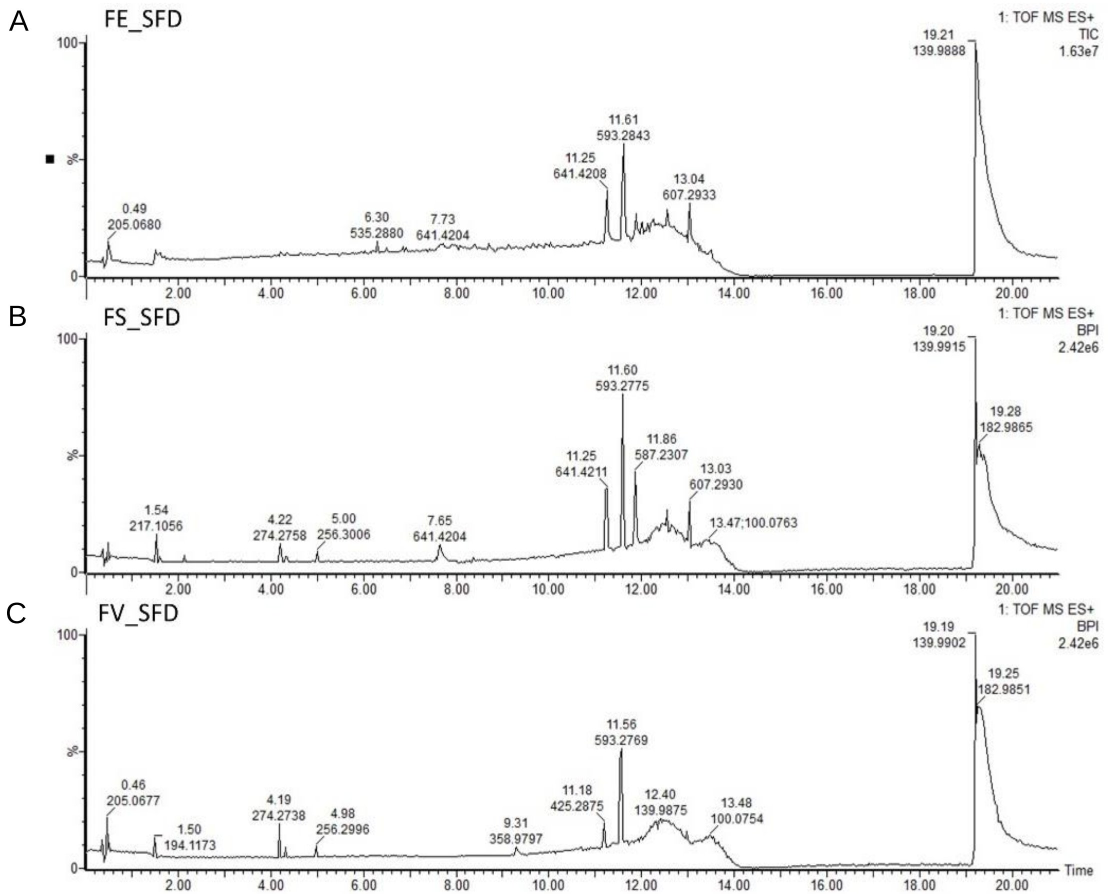


Figure S4: UPLC-MS(+) base peak chromatograms of Surface-free after dipping (SFD) extracts of A) FE, B) FS and C) FV

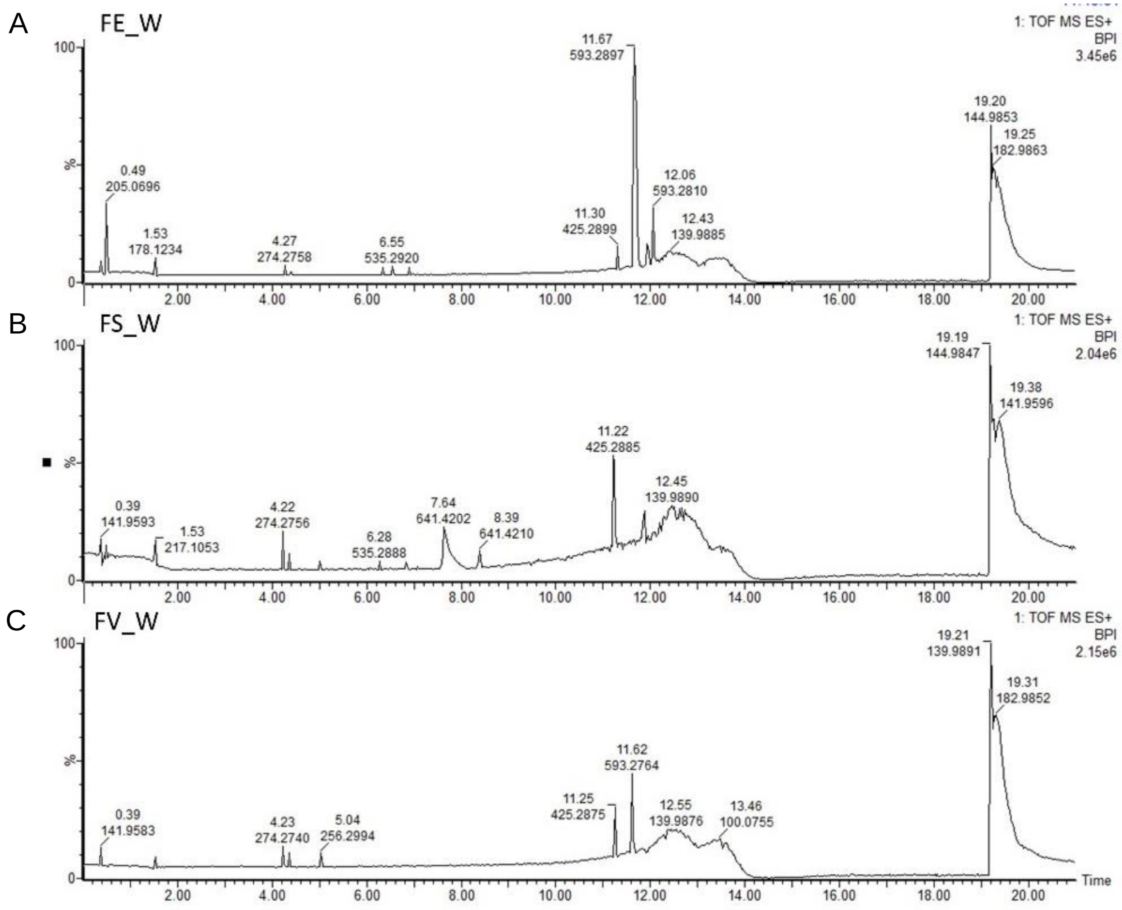


Figure S5: UPLC-MS(+) base peak chromatograms of Whole (W) extracts of A) FE, B) FS and C) FV

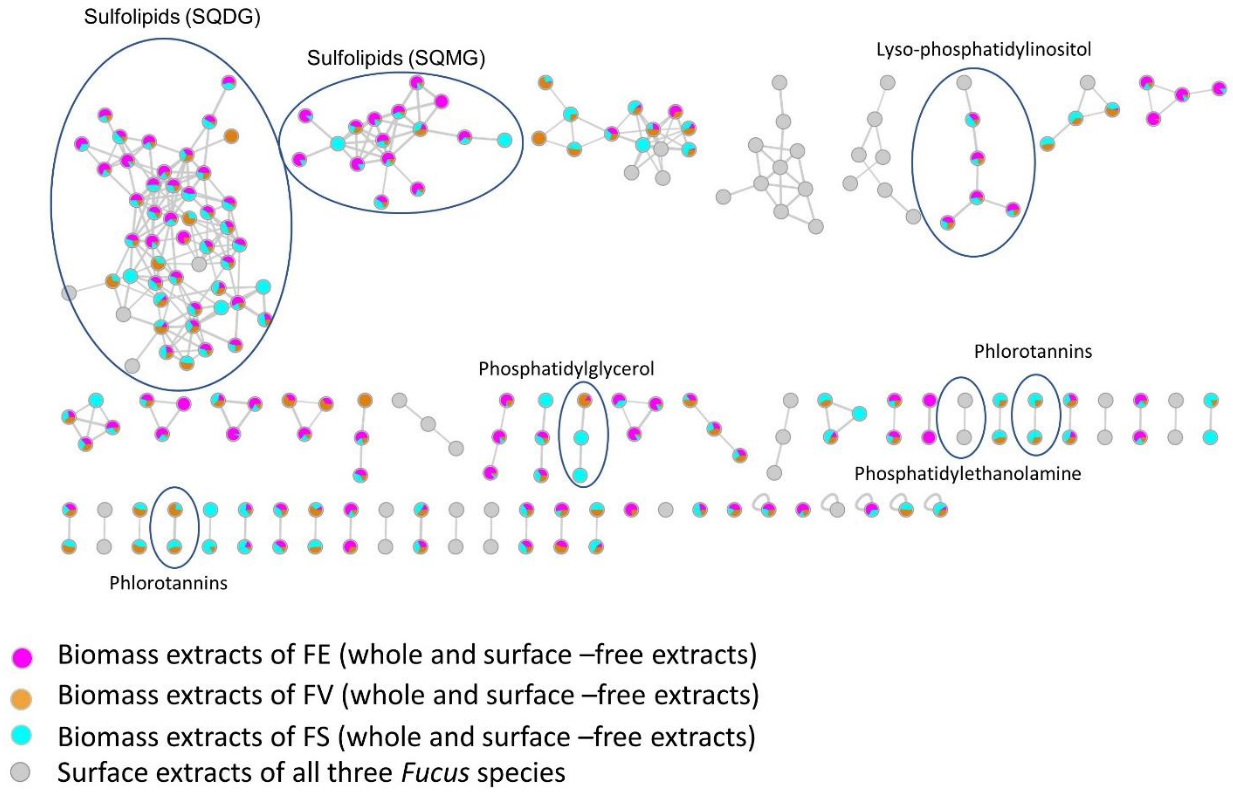


Figure S6: MN generated from UPLC-(-)-ESI-MS/MS data of all extracts of *Fucus* spp. in the negative ion mode. Node colour represents the source of the ion.

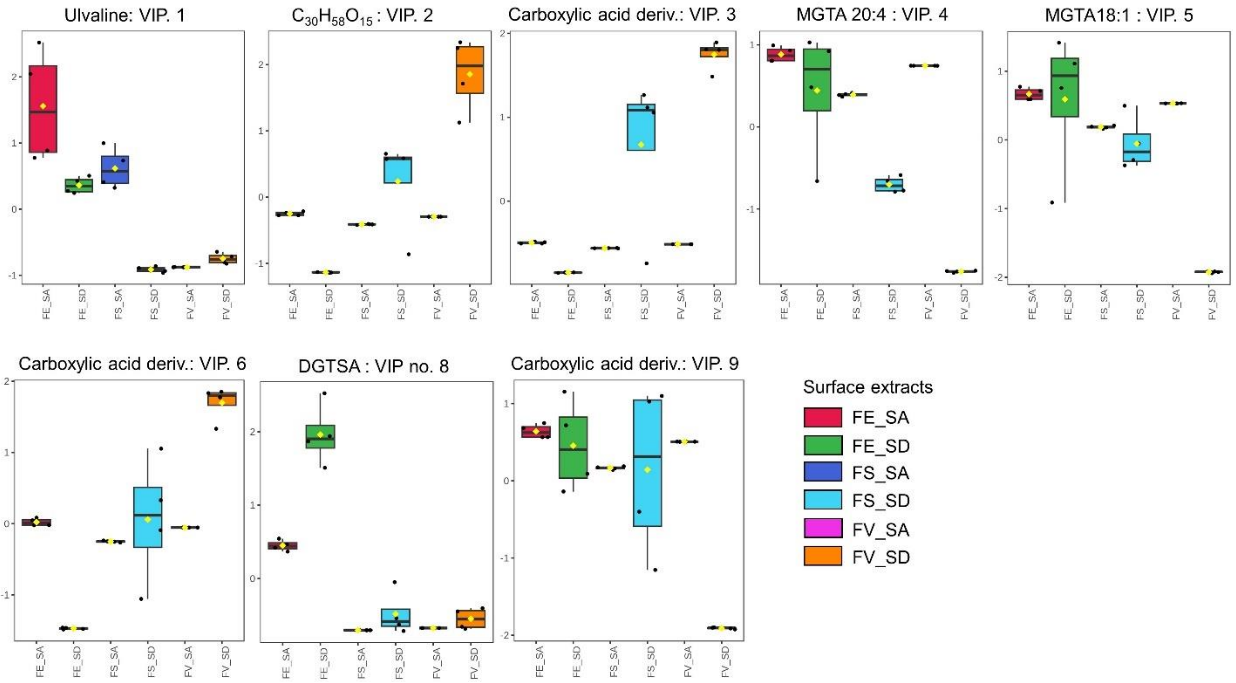


Figure S7. Variation of the most significant discriminatory metabolite markers (VIP >1.8) on the surfaces of the three *Fucus* spp. Normalized intensities are presented on the y-axis.

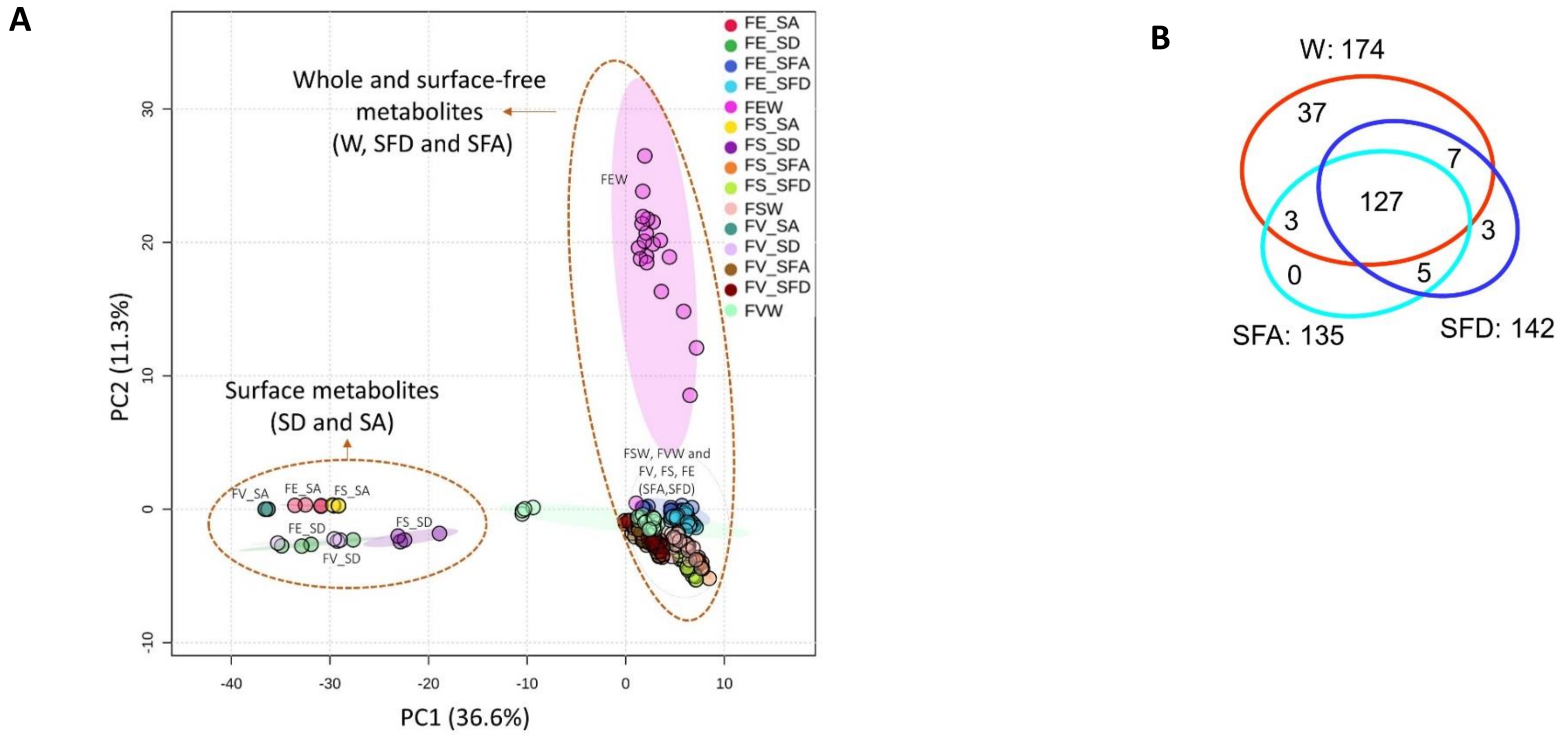


Figure S8: PCA scores plot generated from UPLC-(+)-ESI-MS data of all extracts showing a clear discrimination between surface metabolites from surface free and whole metabolome. Algal species: FV: *F. vesiculosus*, FS: *F. serratus*, FE: *F. distichus* subsp. *evanescens*, Extracts: SA: surface adsorption, SD: solvent dipping, SFA: surface-free after adsorption, SFD: surface-free after dipping, W: whole, untreated algae (B) Venn diagram showing distribution of nodes among the whole extracts (W), surface-free extract after adsorption (SFD) and dipping (SFD).

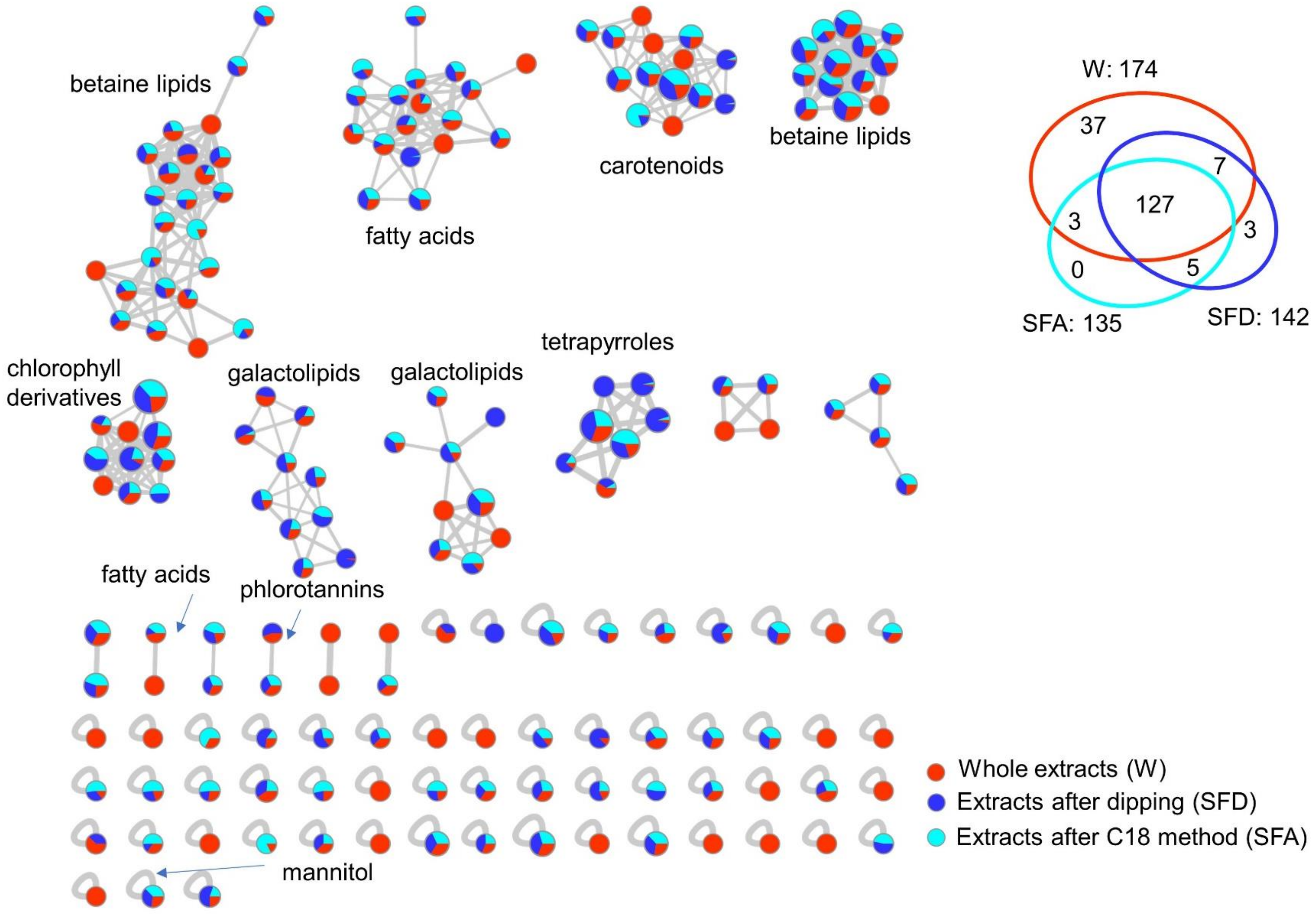
betaine lipids
fatty acids
carotenoids
betaine lipids
W: 174
37
7
127
3
3
0
5
SFA: 135
SFD: 142
chlorophyll derivatives
galactolipids
galactolipids
tetrapyrroles
fatty acids
phlorotannins
mannitol
Whole extracts (W)
Extracts after dipping (SFD)
Extracts after C18 method (SFA)

Figure S9: MN generated from UPLC-(+)-ESI-MS/MS data of all surface-free and whole extracts. Node sizes are modulated according to the sum of intensities of the ions in all extracts while the colors in the pie chart of each node represents the relative quantity of the ion from whole extracts (red), surface-free after dipping (blue) and surface-free after C18 adsorption (light blue)

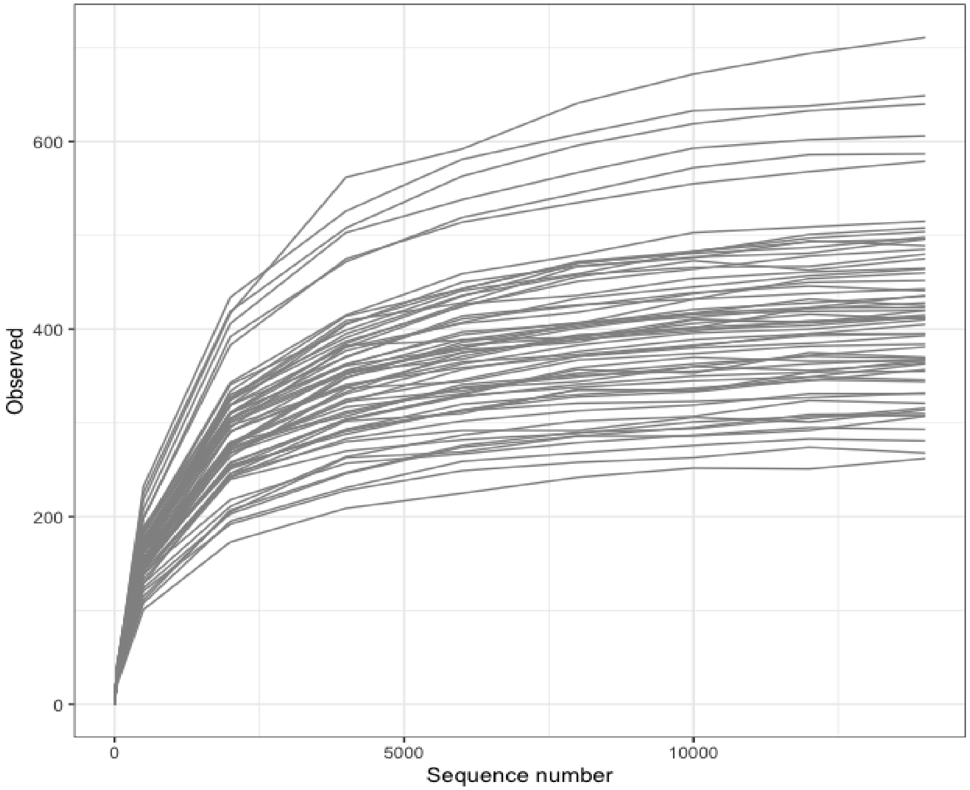


Figure S10: Rarefaction curves of bacterial V3/V4 region amplicon sequences from all 66 samples.

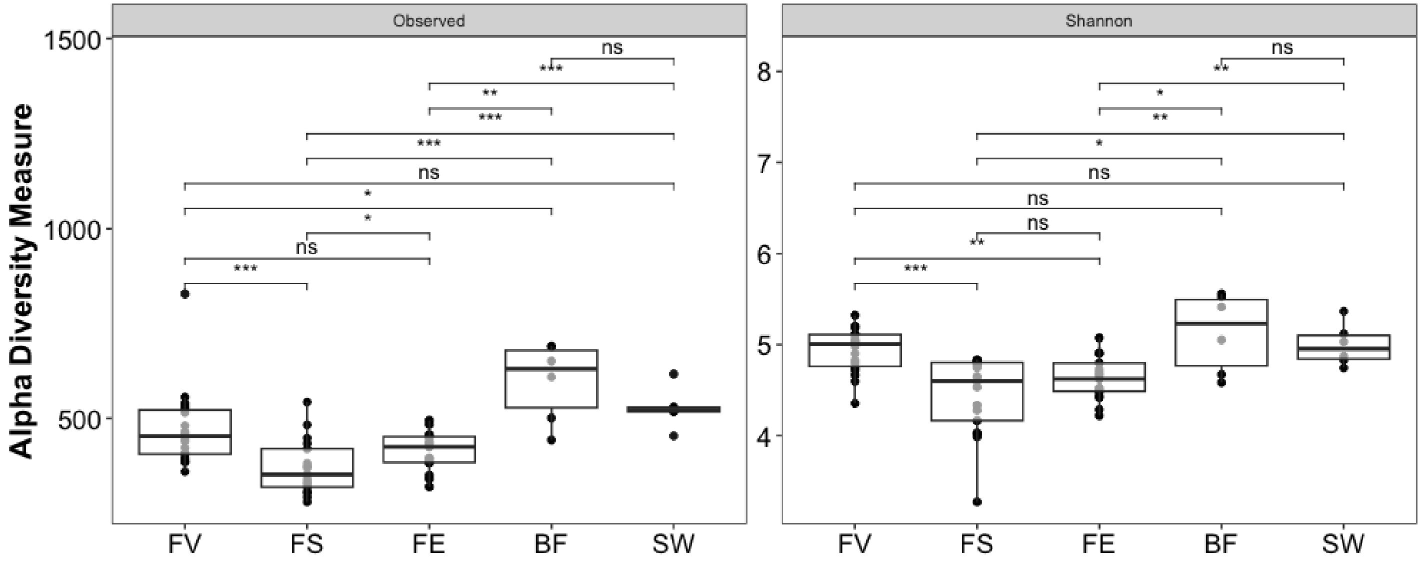


Figure S11: Alpha diversity (ASV-Observed vs. Shannon) of bacterial epiphytic community with regard to sample source. FV: *F. vesiculosus*, FS: *F. serratus*, FE: *F. distichus* subsp. *evanescens,* BF: biofilm on stone, SW: seawater. Significance levels: >0.0001: ***, >0.001: **, 0.01: *, >0.05: ns)

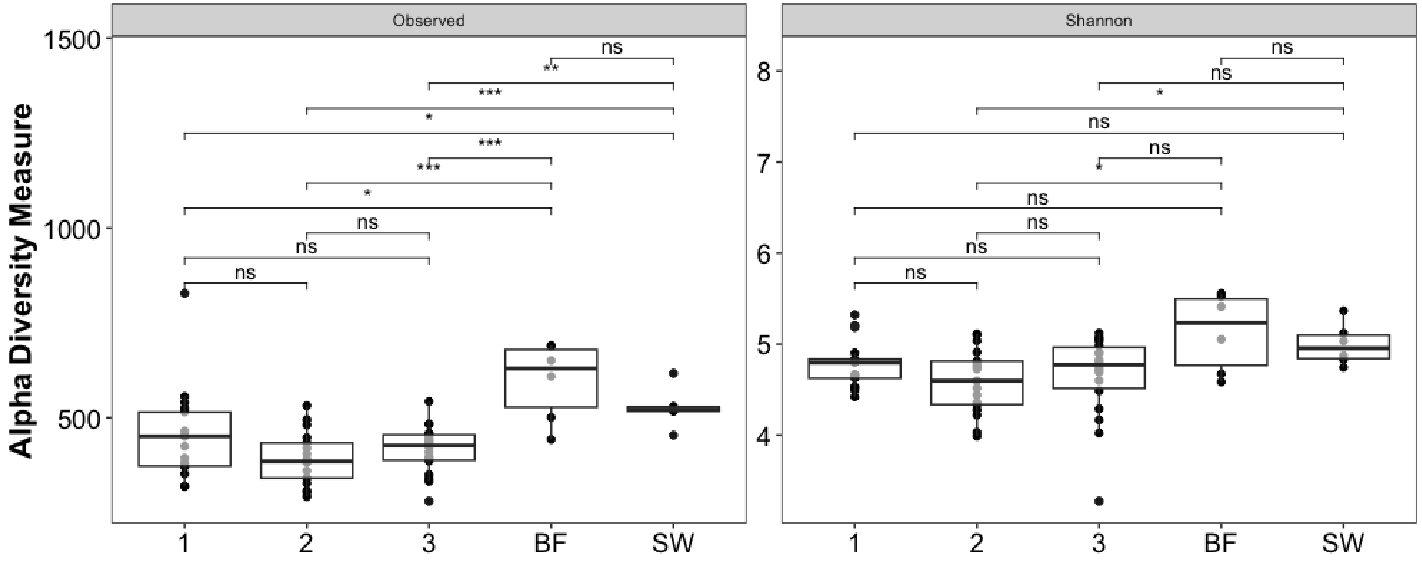


Figure S12: Alpha diversity (ASV-Observed vs. Shannon) of bacterial epiphytic community with regard to individual. 1: individual 1, 2: individual 2, 3: individual 3, BF: biofilm on stone, SW: seawater. Significance levels: >0.0001: ***, >0.001: **, 0.01: *, >0.05: ns)

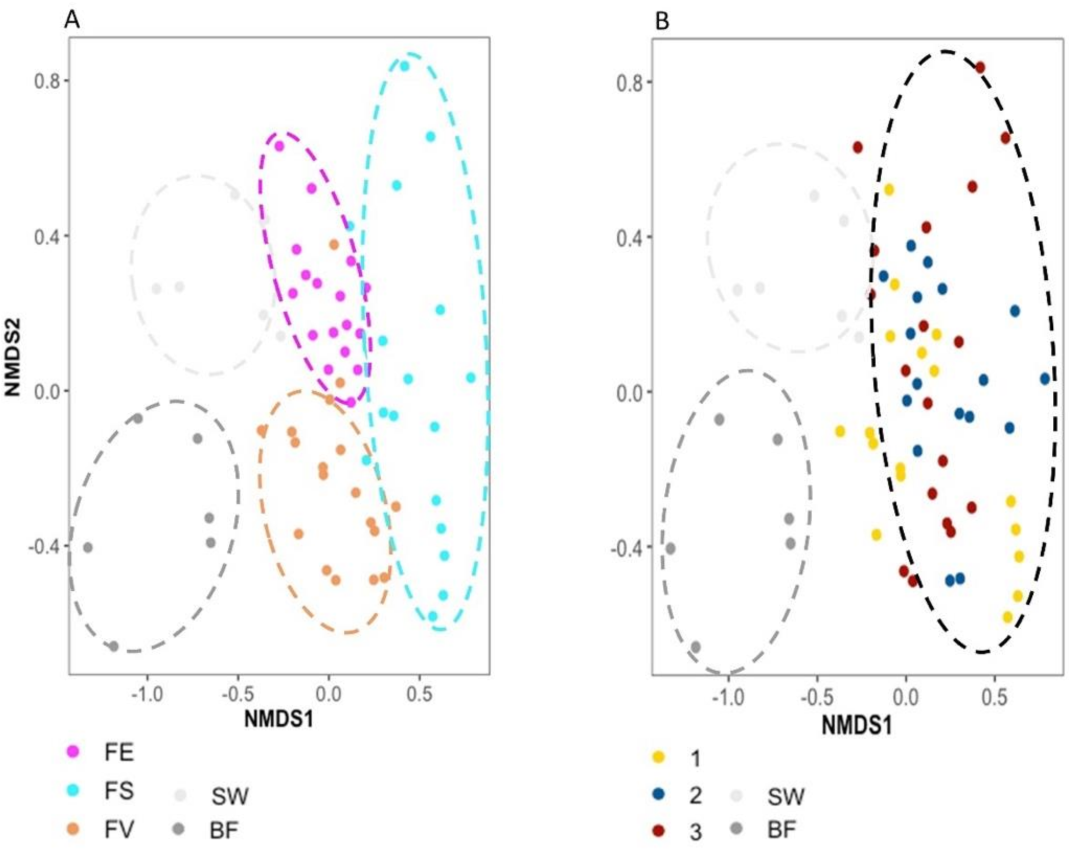


Figure S13: Beta diversity analysis of bacterial amplicon data based on Bray-Curtis distance calculation visualized by NMDS plots (A) according to sample origin (B) according to individual. FV: *Fucus vesiculosus*, FS: *F. serratus,* FE*: F. distichus* subsp. *evanescens,* BF: biofilm on stone, SW: seawater.

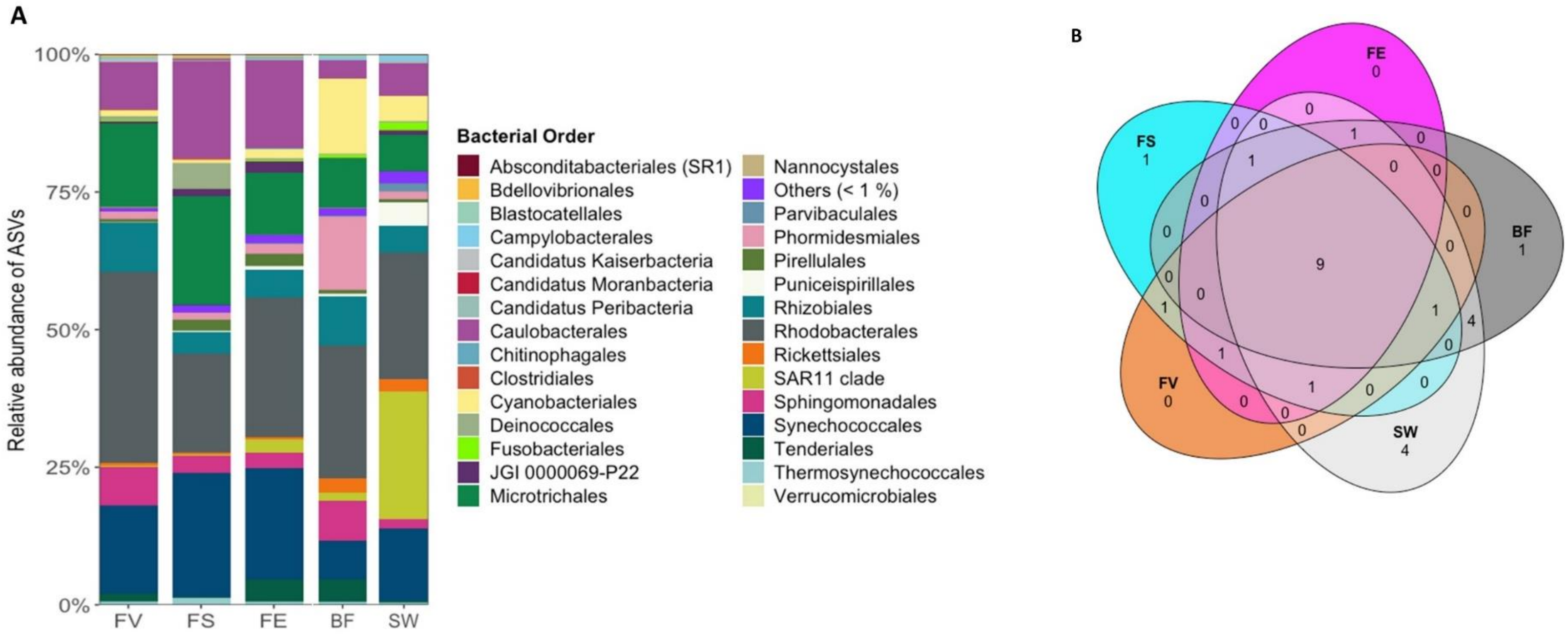


Figure S14: (A) Bacterial orders associated to surfaces of *Fucus* spp., and stone biofilm and seawater reference samples. Others (<1%) represents several orders with less than 1% relative abundance. (B) Venn diagram displaying bacterial orders with regard to different sample types FV: *Fucus vesiculosus*, FS: *F. serratus,* FE*: F. distichus* subsp. *evanescens,* BF: biofilm on stone, SW: seawater.

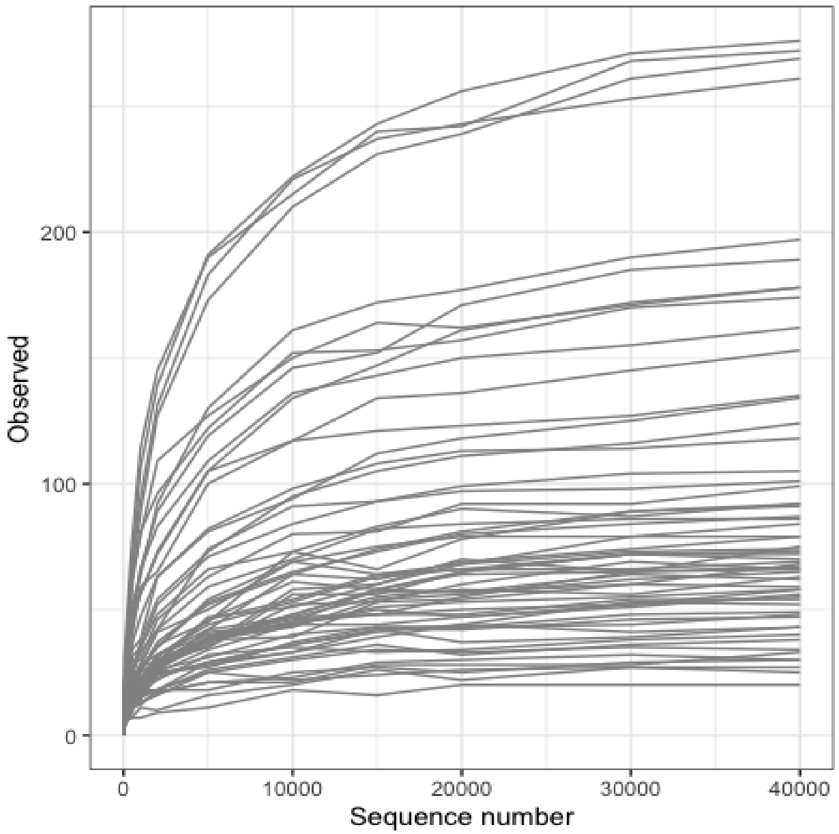


Figure S15: Rarefaction curves of eukaryotic ITS region amplicon sequences from all 60 samples.

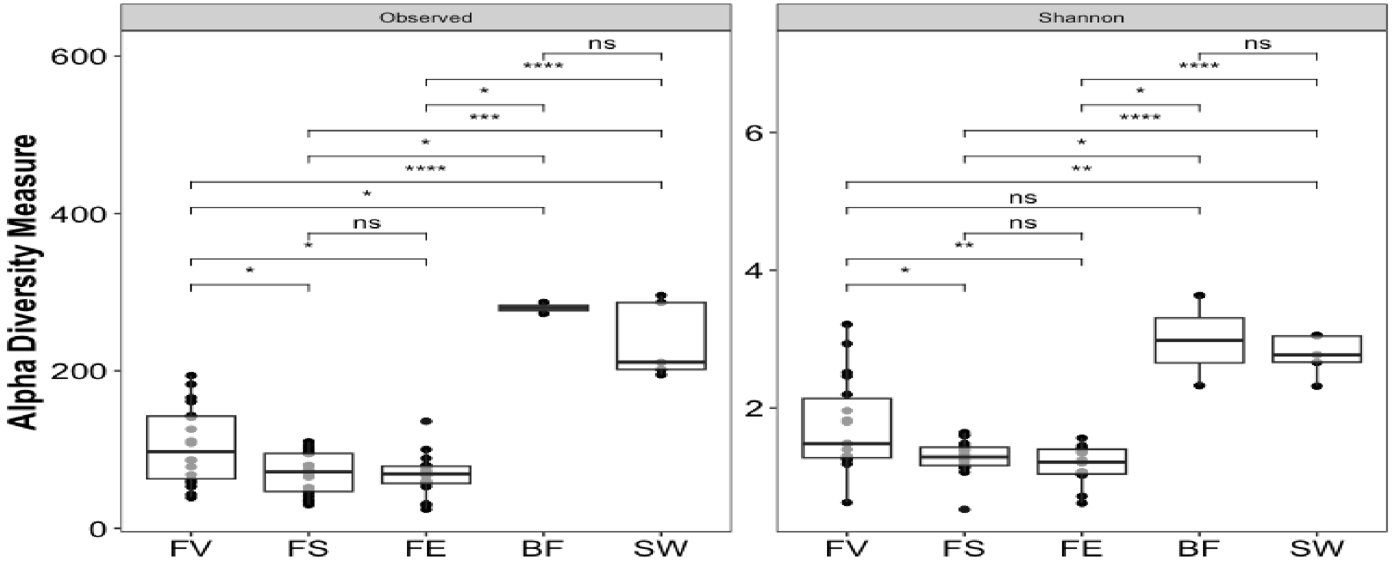


Figure S16: Alpha diversity (ASV-Observed vs. Shannon) of eukaryotic epiphytic community based on ITS fragment sequences with regard to sample source. FV: *F. vesiculosus*, FS: *F. serratus*, FE: *F. distichus* subsp. *evanescens,* BF: biofilm on stone, SW: seawater. Significance levels: >0:****, >0.0001: ***, >0.001: **, 0.01: *, >0.05: ns)

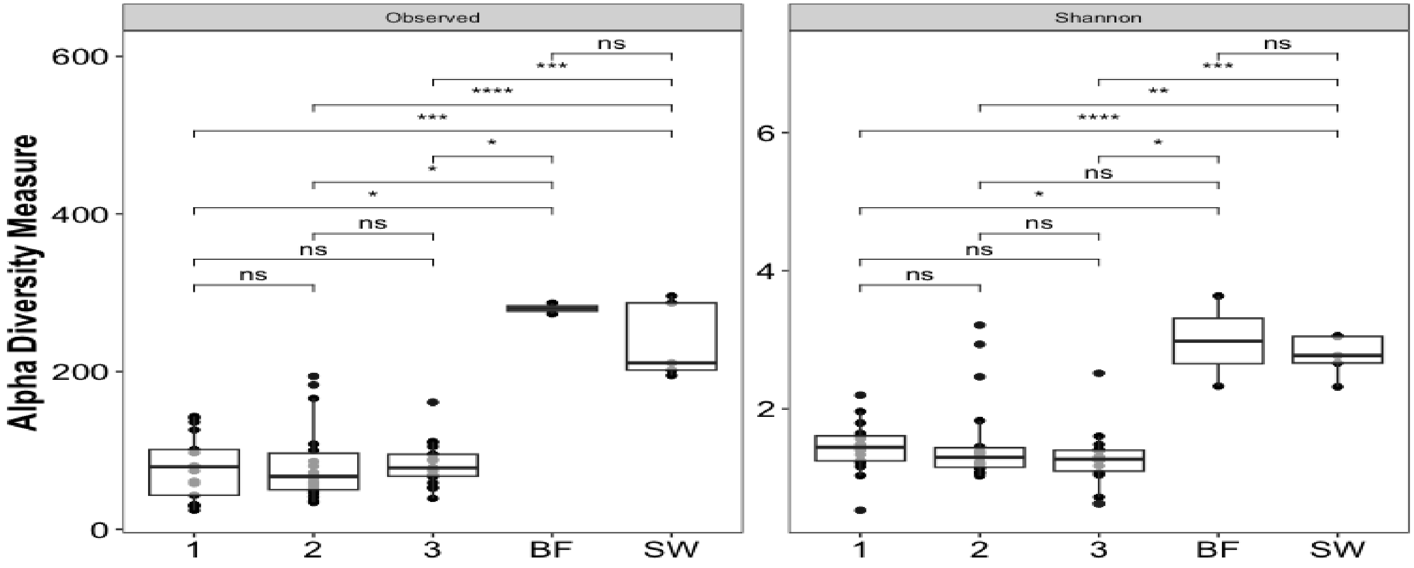


Figure S17: Alpha diversity (ASV-Observed vs. Shannon) of eukaryotic epiphytic community based on ITS fragment sequences with regard to individual. 1: individual 1, 2: individual 2, 3: individual 3, BF: biofilm on stone, SW: seawater. Significance levels: >0:****, >0.0001: ***, >0.001: **, 0.01: *, >0.05: ns)

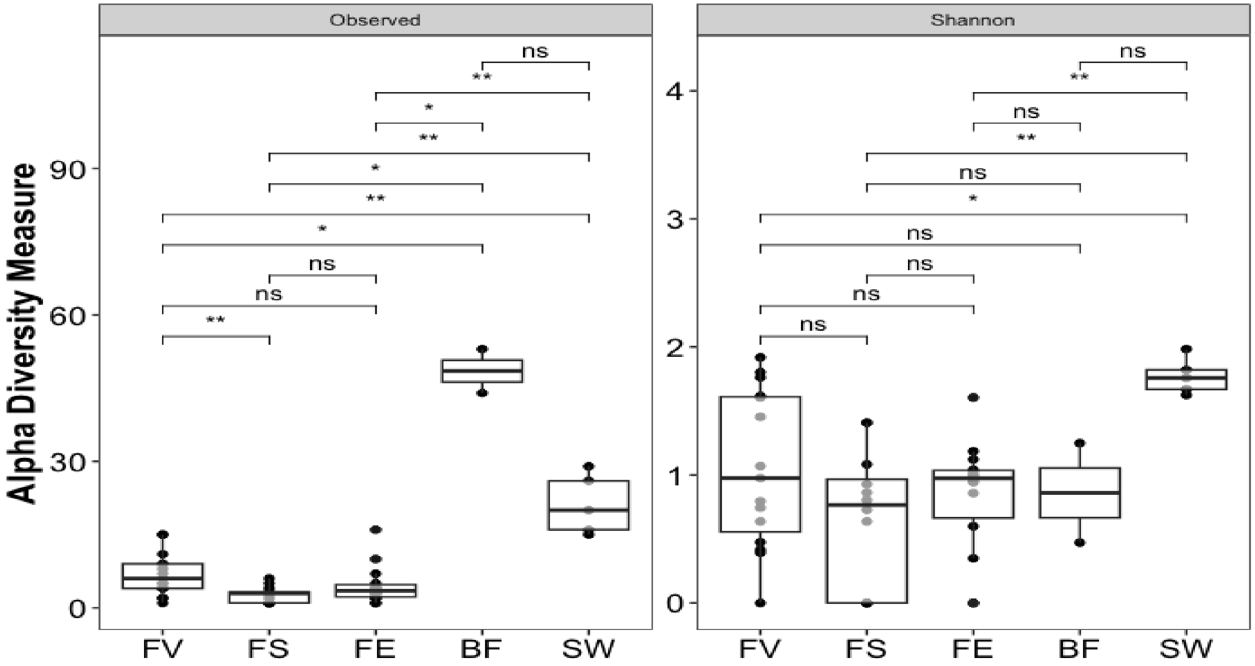


Figure S18: Alpha diversity (ASV-Observed vs. Shannon) of fungal epiphytic community based on ITS fragment sequences with regard to sample source. FV: *F. vesiculosus*, FS: *F. serratus*, FE: *F. distichus* subsp. *evanescens,* BF: biofilm on stone, SW: seawater. Significance levels: >0:****, >0.0001: ***, >0.001: **, 0.01: *, >0.05: ns)

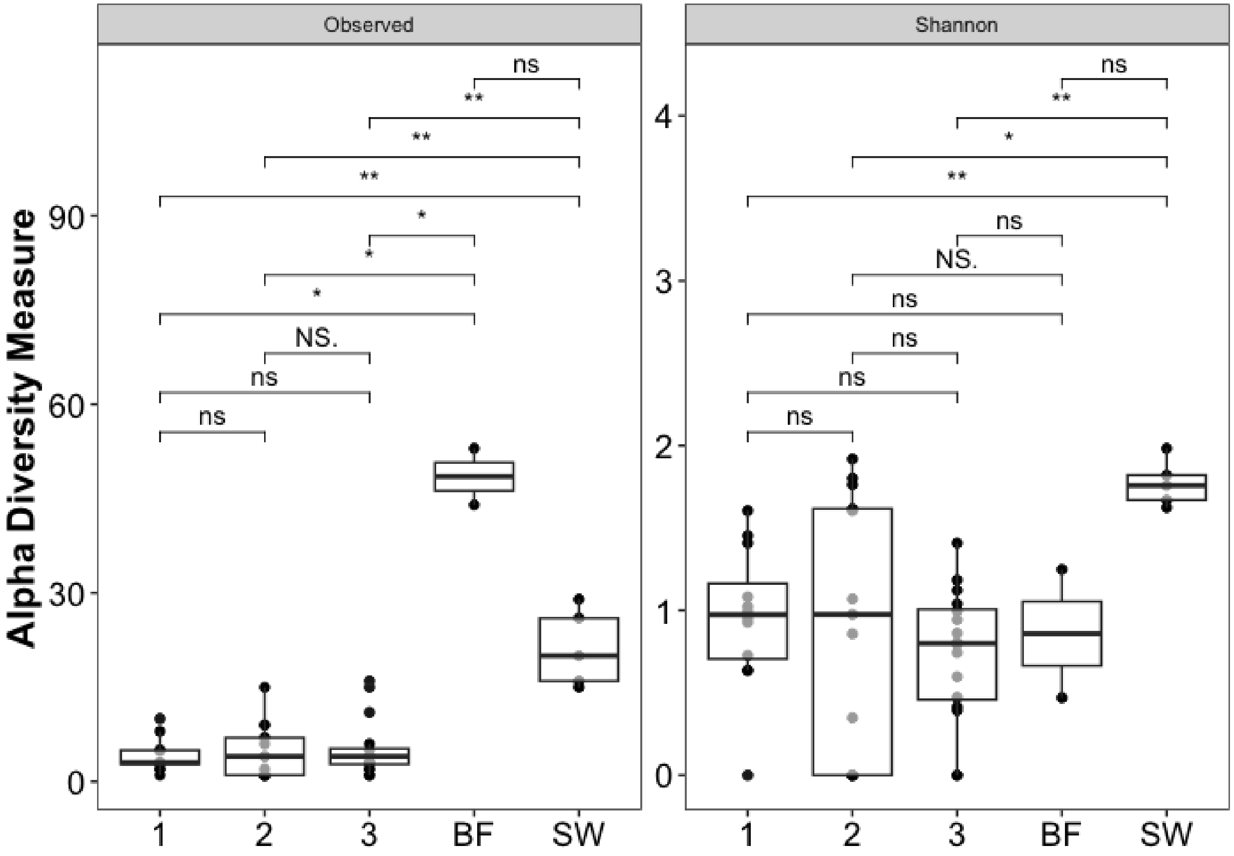


Figure S19: Alpha diversity (ASV-Observed vs. Shannon) of fungal epiphytic community based on ITS fragment sequences with regard to individual. 1: individual 1, 2: individual 2, 3: individual 3, BF: biofilm on stone, SW: seawater. Significance levels: >0:****, >0.0001: ***, >0.001: **, 0.01: *, >0.05: ns)

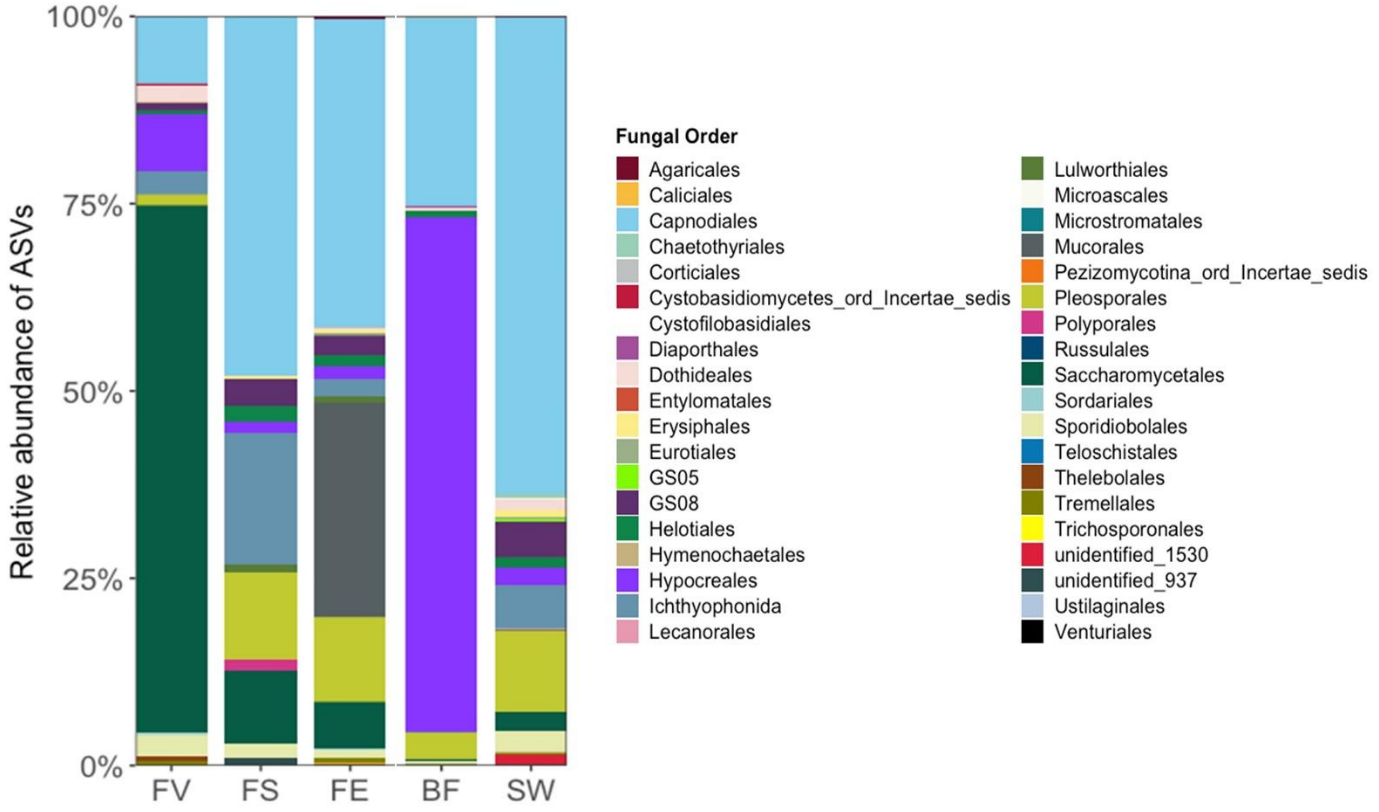


Figure S20: Fungal orders associated to surfaces of *Fucus* spp., and stone biofilm (BF) and seawater (SW) reference samples. FV: *Fucus vesiculosus*, FS: *F. serratus*, FE: *F. distichus* subsp. *evanescens*.

Table S2: Relative abundances of bacterial genera (> 1%) associated to surfaces of *Fucus* spp., seawater and stone biofilm based on amplicon sequencing of the V3/V4 region of the 16S rRNA gene.

| *F. vesiculosus (FV)* | | | | *F. serratus (FS)* | | | | *F. distichus subsp. evanescens (FE)* | | | |
|---|---|---|---|---|---|---|---|---|---|---|---|
| **ASV No.** | **Genus** | **Phylum** | **% Abun-dance** | **ASV No.** | **Genus** | **Phylum** | **% Abun-dance** | **ASV No.** | **Genus** | **Phylum** | **% Abun-dance** |
| ASV1 | Schizothrix LEGE 07164 | Cyanobacteria | 24.41 | ASV1 | Schizothrix LEGE 07164 | Cyanobacteria | 29.61 | ASV1 | Schizothrix LEGE 07164 | Cyanobacteria | 26.02 |
| ASV11 | Sva0996 marine group | Actinobacteriota | 11.78 | ASV11 | Sva0996 marine group | Actinobacteriota | 13.97 | ASV4 | Robiginitomaculum | Proteobacteria | 12.84 |
| ASV12 | Ilumatobacter | Actinobacteriota | 9.85 | ASV17 | Litorimonas | Proteobacteria | 13.65 | ASV10 | Yoonia-Loktanella | Proteobacteria | 8.96 |
| ASV10 | Yoonia-Loktanella | Proteobacteria | 8.68 | ASV12 | Ilumatobacter | Actinobacteriota | 8.31 | ASV11 | Sva0996 marine group | Actinobacteriota | 7.20 |
| ASV20 | Octadecabacter | Proteobacteria | 6.85 | ASV24 | Truepera | Deinococcota | 6.24 | ASV12 | Ilumatobacter | Actinobacteriota | 7.12 |
| ASV17 | Litorimonas | Proteobacteria | 5.08 | ASV4 | Robiginitomaculum | Proteobacteria | 3.62 | ASV13 | Candidatus Tenderia | Proteobacteria | 5.25 |
| ASV14 | Silicimonas | Proteobacteria | 5.00 | ASV21 | Fretibacter | Proteobacteria | 3.50 | ASV20 | Octadecabacter | Proteobacteria | 3.96 |
| ASV51 | Erythrobacter | Proteobacteria | 4.08 | ASV10 | Yoonia-Loktanella | Proteobacteria | 2.99 | ASV17 | Litorimonas | Proteobacteria | 3.86 |
| ASV4 | Robiginitomaculum | Proteobacteria | 2.74 | ASV68 | Blastopirellula | Planctomycetota | 2.67 | ASV68 | Blastopirellula | Planctomycetota | 2.90 |
| ASV39 | Filomicrobium | Proteobacteria | 2.48 | ASV20 | Octadecabacter | Proteobacteria | 2.36 | ASV21 | Fretibacter | Proteobacteria | 2.69 |
| ASV13 | Candidatus Tenderia | Proteobacteria | 2.04 | ASV88 | Acaryochloris MBIC11017 | Cyanobacteria | 1.71 | ASV19 | Clade Ia | Proteobacteria | 2.03 |
| ASV73 | Altererythrobacter | Proteobacteria | 1.54 | ASV14 | Silicimonas | Proteobacteria | 1.59 | ASV14 | Silicimonas | Proteobacteria | 1.75 |
| ASV21 | Fretibacter | Proteobacteria | 1.35 | ASV61 | Phormidesmis ANT.LACV5.1 | Cyanobacteria | 1.39 | ASV61 | Phormidesmis ANT.LACV5.1 | Cyanobacteria | 1.64 |
| ASV61 | Phormidesmis ANT.LACV5.1 | Cyanobacteria | 1.24 | ASV39 | Filomicrobium | Proteobacteria | 1.05 | ASV29 | Planktomarina | Proteobacteria | 1.31 |
| ASV24 | Truepera | Deinococcota | 1.09 | | | | | | | | |

| *seawater (SW)* | | | | *stone biofilm (BF)* | | | |
|---|---|---|---|---|---|---|---|
| **ASV No.** | **Genus** | **Phylum** | **% Abun-dance** | **ASV No.** | **Genus** | **Phylum** | **% Abun-dance** |
| ASV19 | Clade Ia | Proteobacteria | 22.46 | ASV61 | Phormidesmis ANT.LACV5.1 | Cyanobacteria | 12.76 |
| ASV1 | Schizothrix LEGE 07164 | Cyanobacteria | 14.56 | ASV49 | Rivularia PCC-7116 | Cyanobacteria | 11.30 |
| ASV29 | Planktomarina | Proteobacteria | 10.17 | ASV12 | Ilumatobacter | Actinobacteriota | 9.39 |
| ASV79 | Cand. Puniceispirillum | Proteobacteria | 6.02 | ASV1 | Schizothrix LEGE 07164 | Cyanobacteria | 8.85 |
| ASV12 | Ilumatobacter | Actinobacteriota | 4.89 | ASV13 | Cand. Tenderia | Proteobacteria | 5.80 |
| ASV11 | Sva0996 marine group | Actinobacteriota | 4.13 | ASV10 | Yoonia-Loktanella | Proteobacteria | 5.07 |
| ASV4 | Robiginitomaculum | Proteobacteria | 4.09 | ASV76 | Sphingorhabdus | Proteobacteria | 4.66 |
| ASV130 | Aphanizomenon NIES81 | Cyanobacteria | 3.44 | ASV116 | Pleurocapsa PCC-7319 | Cyanobacteria | 4.37 |
| ASV158 | Cyanobium PCC-6307 | Cyanobacteria | 3.42 | ASV93 | Phormidium MBIC10003 | Cyanobacteria | 4.05 |
| ASV10 | Yoonia-Loktanella | Proteobacteria | 3.38 | ASV11 | Sva0996 marine group | Actinobacteriota | 3.06 |
| ASV171 | Propionigenium | Fusobacteriota | 2.09 | ASV51 | Erythrobacter | Proteobacteria | 1.89 |
| ASV17 | Litorimonas | Proteobacteria | 1.88 | ASV98 | Ahrensia | Proteobacteria | 1.61 |
| ASV20 | Octadecabacter | Proteobacteria | 1.73 | ASV4 | Robiginitomaculum | Proteobacteria | 1.54 |
| ASV61 | Phormidesmis ANT.LACV5.1 | Cyanobacteria | 1.52 | ASV14 | Silicimonas | Proteobacteria | 1.34 |
| ASV14 | Silicimonas | Proteobacteria | 1.28 | ASV19 | Clade Ia | Proteobacteria | 1.25 |
| | | | | ASV158 | Cyanobium PCC-6307 | Cyanobacteria | 1.22 |
| | | | | ASV29 | Planktomarina | Proteobacteria | 1.12 |
| | | | | ASV20 | Octadecabacter | Proteobacteria | 1.10 |

Table S3: Bacterial beta diversity statistics based on Bray-Curtis dissimilarity, left: PERMANOVA results, right: pairwise permutation test for homogeneity of multivariate dispersions

| PERMANOVA | | Signif. codes: 0 ‘***’ 0.001 ‘**’ 0.01 ‘*’ 0.05 ‘.’ 0.1 ‘ ’ 1 | | | | | | | | pairwise permutation for homogeneity of multivariate dispersion | | | | | | | | | |
|---|---|---|---|---|---|---|---|---|---|---|---|---|---|---|---|---|---|---|---|
| | Df | Sum Squares | R2 | F | Pr(>F) | Signifi-cance | Resi-duals df | Residuals Sum Squares | Residuals R2 | Df | Sum Squares | Mean Squares | F | N permut | Pr(>F) | Signifi-cance | Resi-duals df | Residuals sum squares | Residuals mean squares |
| **full dataset** | | | | | | | | | | **full dataset** | | | | | | | | | |
| origin | 4 | 4.8181 | 0.39588 | 9.6655 | 0.001 | *** | 59 | 7.3572 | 0.60412 | 4 | 0.13976 | 0.034941 | 8.1559 | 1000 | 0.000999 | *** | 59 | 0.25276 | 0.004284 |
| indivdual no. | 4 | 3.1344 | 0.25753 | 5.1162 | 0.001 | *** | 59 | 9.0364 | 0.74247 | 4 | 0.13753 | 0.034383 | 7.4047 | 1000 | 0.000999 | *** | 59 | 0.27396 | 0.004643 |
| **FV** | | | | | | | | | | **FV** | | | | | | | | | |
| indivdual no. | 2 | 0.50452 | 0.26094 | 2.6481 | 0.015 | * | 15 | 1.42895 | 0.73906 | 2 | 0.00199 | 0.000966 | 0.2325 | 1000 | 0.7972 | ns | 15 | 0.064251 | 0.004283 |
| tissue age | 1 | 0.58970 | 0.30499 | 7.0214 | 0.001 | *** | 16 | 1.3438 | 0.69501 | 1 | 0.00586 | 0.005862 | 1.361 | 1000 | 0.2677 | ns | 16 | 0.068909 | 0.004307 |
| **FS** | | | | | | | | | | **FS** | | | | | | | | | |
| indivdual no. | 2 | 1.16040 | 0.42611 | 5.1974 | 0.001 | *** | 14 | 1.5629 | 0.57389 | 2 | 0.03316 | 0.01658 | 2.1023 | 1000 | 0.1638 | ns | 14 | 0.110414 | 0.007887 |
| **FE** | | | | | | | | | | **FE** | | | | | | | | | |
| indivdual no. | 2 | 0.39425 | 0.23931 | 2.2022 | 0.013 | * | 14 | 1.25319 | 0.76069 | 2 | 0.00709 | 0.003544 | 1.0939 | 1000 | 0.3716 | ns | 14 | 0.045354 | 0.00324 |

Table S4: Relative abundances of eukaryote genera (> 1%) associated to surfaces of *Fucus* spp., seawater and stone biofilm based on amplicon sequencing of the ITS fragment.

| *F. vesiculosus* (FV) | | | |
|---|---|---|---|
| **ASV No.** | **Genus** | **Phylum** | **% Abundance** |
| ASV52 | Zoothamnium | Ciliophora | 39.26 |
| ASV45 | unidentified_91524 | Ciliophora | 28.84 |
| ASV64 | Candida | Ascomycota | 16.26 |
| ASV106 | Pseudovorticella | Ciliophora | 3.83 |
| ASV178 | unidentified_15 | unidentified | 2.32 |
| ASV210 | Myoschiston | Ciliophora | 1.92 |
| ASV10 | Haptocillium | Ascomycota | 1.82 |
| ASV216 | unidentified_4835 | unidentified_6334 | 1.62 |

| *F. serratus* (FS) | | | |
|---|---|---|---|
| **ASV No.** | **Genus** | **Phylum** | **% Abundance** |
| ASV45 | unidentified_91524 | Ciliophora | 33.17 |
| ASV87 | Ulva | Chlorophyta | 17.18 |
| ASV19 | unidentified_12446 | Ciliophora | 11.57 |
| ASV106 | Pseudovorticella | Ciliophora | 9.69 |
| ASV28 | Sarsia | Cnidaria | 7.14 |
| ASV280 | Uronema_91786 | Ciliophora | 5.94 |
| ASV108 | unidentified_5383 | unidentified_6967 | 4.17 |
| ASV366 | Haliclystus | Cnidaria | 2.07 |
| ASV52 | Zoothamnium | Ciliophora | 1.25 |

| *F. distichus* subsp. *evanescens* (FE) | | | |
|---|---|---|---|
| **ASV No.** | **Genus** | **Phylum** | **% Abundance** |
| ASV28 | Sarsia | Cnidaria | 27.22 |
| ASV19 | unidentified_12446 | Ciliophora | 25.46 |
| ASV221 | Mucor | Mucoromycota | 11.60 |
| ASV87 | Ulva | Chlorophyta | 8.74 |
| ASV45 | unidentified_91524 | Ciliophora | 6.17 |
| ASV108 | unidentified_5383 | unidentified_6967 | 3.81 |
| ASV97 | Alternaria | Ascomycota | 2.64 |
| ASV127 | Pseudendoclonium | Chlorophyta | 2.22 |
| ASV106 | Pseudovorticella | Ciliophora | 2.08 |
| ASV166 | unidentified_33279 | Chlorophyta | 1.23 |
| ASV244 | unidentified_2488 | Rozellomycota | 1.02 |

| seawater (SW) | | | |
|---|---|---|---|
| **ASV No.** | **Genus** | **Phylum** | **% Abundance** |
| ASV28 | Sarsia | Cnidaria | 39.62 |
| ASV19 | unidentified_12446 | Ciliophora | 35.05 |
| ASV108 | unidentified_5383 | unidentified_6967 | 5.81 |
| ASV45 | unidentified_91524 | Ciliophora | 4.89 |
| ASV87 | Ulva | Chlorophyta | 3.30 |
| ASV106 | Pseudovorticella | Ciliophora | 1.94 |

| stone biofilm (BF) | | | |
|---|---|---|---|
| **ASV No.** | **Genus** | **Phylum** | **% Abundance** |
| ASV10 | Haptocillium | Ascomycota | 76.14 |
| ASV87 | Ulva | Chlorophyta | 5.84 |
| ASV75 | Cladosporium | Ascomycota | 3.94 |
| ASV127 | Pseudendoclonium | Chlorophyta | 3.47 |
| ASV97 | Alternaria | Ascomycota | 2.80 |
| ASV45 | unidentified_91524 | Ciliophora | 2.34 |
| ASV166 | unidentified_33279 | Chlorophyta | 1.17 |

Table S5: ITS beta diversity statistics based on Bray-Curtis dissimilarity, left: PERMANOVA results, right: pairwise permutation test for homogeneity of multivariate dispersions

**PERMANOVA** — Signif. codes: 0 ‘***’ 0.001 ‘**’ 0.01 ‘*’ 0.05 ‘.’ 0.1 ‘ ’ 1

| | Df | Sum Squares | R2 | F | Pr(>F) | Signifi-cance | Resi-duals df | Residuals Sum Squares | Residuals R2 |
|---|---|---|---|---|---|---|---|---|---|
| **full dataset** | | | | | | | | | |
| origin | 4 | 4.0572 | 0.29338 | 5.7089 | 0.001 | *** | 55 | 9.7718 | 0.70662 |
| indivdual no. | 4 | 3.7628 | 0.2721 | 5.1398 | 0.001 | *** | 55 | 10.0662 | 0.7279 |
| **FV** | | | | | | | | | |
| indivdual no. | 2 | 0.7918 | 0.2223 | 2.1438 | 0.054 | . | 15 | 2.77 | 0.7777 |
| **FS** | | | | | | | | | |
| indivdual no. | 2 | 0.37763 | 0.18583 | 1.7118 | 0.076 | . | 15 | 1.65453 | 0.81417 |
| **FE** | | | | | | | | | |
| indivdual no. | 2 | 0.27035 | 0.09155 | 0.7054 | 0.493 | ns | 14 | 2.68281 | 0.90845 |

**pairwise permutation for homogeneity of multivariate dispersion**

| | Df | Sum Squares | Mean Squares | F | N permut | Pr(>F) | Signifi-cance | Resi-duals df | Residuals sum squares | Residuals mean squares |
|---|---|---|---|---|---|---|---|---|---|---|
| **full dataset** | | | | | | | | | | |
| origin | 4 | 0.13976 | 0.034941 | 8.1559 | 1000 | 0.000999 | *** | 59 | 0.25276 | 0.004284 |
| indivdual no. | 4 | 0.13753 | 0.034383 | 7.4047 | 1000 | 0.000999 | *** | 59 | 0.27396 | 0.004643 |
| **FV** | | | | | | | | | | |
| indivdual no. | 2 | 0.00199 | 0.000966 | 0.2325 | 1000 | 0.7972 | ns | 15 | 0.064251 | 0.004283 |
| **FS** | | | | | | | | | | |
| indivdual no. | 2 | 0.03316 | 0.01658 | 2.1023 | 1000 | 0.1638 | ns | 14 | 0.110414 | 0.007887 |
| **FE** | | | | | | | | | | |
| indivdual no. | 2 | 0.00709 | 0.003544 | 1.0939 | 1000 | 0.3716 | ns | 14 | 0.045354 | 0.00324 |

Table S6: Relative abundances of fungal genera (>1%) associated to the surfaces of *Fucus* spp., seawater and stone biofilm based on amplicon sequencing of the ITS fragment.

| *F. vesiculosus (FV)* | | | | *F. serratus (FS)* | | | | *F. distichus subsp. evanescens (FE)* | | | |
|---|---|---|---|---|---|---|---|---|---|---|---|
| **ASV No.** | **Genus** | **Phylum** | **% Abun-dance** | **ASV No.** | **Genus** | **Phylum** | **% Abun-dance** | **ASV No.** | **Genus** | **Phylum** | **% Abun-dance** |
| ASV64 | Candida | Ascomycota | 82.30 | ASV186 | Sphaeroforma | Ichthyosporia_phy_Incertae_sedis | 38.04 | ASV221 | Mucor | Mucoromycota | 65.41 |
| ASV10 | Haptocillium | Ascomycota | 9.20 | ASV97 | Alternaria | Ascomycota | 25.00 | ASV97 | Alternaria | Ascomycota | 14.91 |
| ASV186 | Sphaeroforma | Ichthyosporia_phy_Incertae_sedis | 3.68 | ASV342 | Metschnikowia | Ascomycota | 17.39 | ASV244 | unidentified_2488 | Rozellomycota | 5.77 |
| ASV97 | Alternaria | Ascomycota | 1.61 | ASV244 | unidentified_2488 | Rozellomycota | 7.61 | ASV186 | Sphaeroforma | Ichthyosporia_phy_Incertae_sedis | 5.37 |
| ASV244 | unidentified_2488 | Rozellomycota | 1.18 | ASV1291 | Wickerhamomyces | Ascomycota | 3.26 | ASV871 | unidentified_99109 | Ascomycota | 2.19 |
| | | | | ASV451 | Claviceps | Ascomycota | 3.26 | | | | |
| | | | | ASV1208 | unidentified_786 | Ascomycota | 2.17 | | | | |
| | | | | ASV871 | unidentified_99109 | Ascomycota | 2.17 | | | | |
| | | | | ASV1107 | Blumeria | Ascomycota | 1.09 | | | | |

| *seawater (SW)* | | | | *stone biofilm (BF)* | | | |
|---|---|---|---|---|---|---|---|
| **ASV No.** | **Genus** | **Phylum** | **% Abun-dance** | **ASV No.** | **Genus** | **Phylum** | **% Abun-dance** |
| ASV97 | Alternaria | Ascomycota | 26.17 | ASV10 | Haptocillium | Ascomycota | 90.82 |
| ASV186 | Sphaeroforma | Ichthyosporia_phy_Incertae_sedis | 17.73 | ASV75 | Cladosporium | Ascomycota | 4.70 |
| ASV244 | unidentified_2488 | Rozellomycota | 14.69 | ASV97 | Alternaria | Ascomycota | 3.35 |
| ASV75 | Cladosporium | Ascomycota | 13.98 | | | | |
| ASV342 | Metschnikowia | Ascomycota | 7.11 | | | | |
| ASV451 | Claviceps | Ascomycota | 4.30 | | | | |
| ASV440 | unidentified_1163 | Chytridiomycota | 3.67 | | | | |
| ASV573 | Pyrenophora | Ascomycota | 2.89 | | | | |
| ASV568 | Knufia | Ascomycota | 1.64 | | | | |
